# Expanded Expression of Pro-Neurogenic Factor SoxB1 during Larval Development of Gastropod *Lymnaea stagnalis* Suggests Preadaptation to Prolonged Neurogenesis in Mollusca

**DOI:** 10.1101/2023.11.29.569209

**Authors:** Anastasia I. Kurtova, Alexander D. Finoshin, Margarita S. Aparina, Guzel R. Gazizova, Olga S. Kozlova, Svetlana N. Voronova, Elena I. Shagimardanova, Evgeny G. Ivashkin, Elena E. Voronezhskaya

**Author notes:** Correspondence: Evgeny Ivashkin Elena Voronezhskaya.

## Abstract

The diversity in the organization of the nervous system in mollusks raises intriguing questions about its development and evolution. Our study aims to gain a deeper understanding of how the nervous system forms in Mollusca by examining the involvement of SoxB-family transcription factors in the early development of neurogenic zones. Specifically, we explore the expression patterns of two SoxB genes in the gastropod *Lymnaea stagnalis*, namely Ls-SoxB1 and Ls-SoxB2, across various developmental stages. Through a combination of in situ hybridization chain reaction, immunohistochemistry, and proliferation assays, we examine the dynamic spatial distribution of Ls- SoxB1 and Ls-SoxB2, with a particular emphasis on the formation of central ring ganglia and the identification of active proliferative zones. Our findings reveal that Ls-SoxB1 exhibits expanded ectodermal expression from the gastrula to the postmetamorphic stage, evident at both transcriptional and translational levels. Throughout larval development, Ls-SoxB1 is expressed in the ectoderm of the head, foot, and visceral complex, as well as in ganglia anlagen and sensory cells. In contrast, the expression of Ls-SoxB2 in the ectoderm is observed until the veliger stage, after which it persists in subepithelial layer cells and ganglia rudiments. Proliferation assay reveals a uniform distribution of dividing cells in the ectoderm at all developmental stages, indicating the absence of distinct neurogenic zones with increased proliferation in gastropods. Our findings highlight that Ls-SoxB1 exhibit widespread expression patterns in both location and time compared to other Lophotrochozoa species. This prolonged expression of SoxB genes in gastropods can be interpreted as a form of transcriptional neoteny, playing a crucial role in the diversification of nervous systems. Thus, it serves as a preadaptation to prolonged neurogenesis and an increase in the central nervous system complexity in Mollusca.

## 1 Introduction

Mollusca, a highly diverse phylum within Bilateria, exhibits an astonishing array of body forms, lifestyles, and ecological adaptations. This remarkable diversity is not only visible in their external morphology but is also deeply intertwined with the developmental patterns of the nervous system of these organisms. The structure of the molluscan nervous system reflects their lifestyles and demonstrates high plasticity at the morphological level, from the scattered ganglia in Bivalves to the highly centralized brain of Cephalopods (Bullock and Horridge, 1965; Schmidt-Rhaesa et al., 2015). And it is an intriguing question how such a diversity of nervous system arises during development and emerges in evolution.

Our knowledge about the formation of the nervous system in mollusks is mostly restricted to the appearance of already differentiated neurons (Croll, 2009; Nielsen, 2012; Richter et al., 2015; Voronezhskaya and Croll, 2015). Spatial and temporal distribution of neurons expressing specific transmitter phenotypes vary significantly between molluscan classes and even within one family, making it difficult for comparative analysis of neurogenesis (Sakharov, 1976; Croll, 2000; Moroz, 2009, 2021). Based on the existing data for other invertebrate groups, it seems reasonable to look at the early neurogenic events. Particularly, the data about neurogenic stem cells and expression of transcriptional factors that prerequisite neuron specification. Such factors have been identified for all the main groups of Neuralia representatives from cnidarians to vertebrates and demonstrated high evolutionarily conserved features (Marlow et al., 2014; Moroz, 2015; Arendt et al., 2016; Arendt, 2018; Martín-Durán et al., 2018). In the case of mollusks, such sets of early and late neurogenic factors have been described in detail only for cephalopods (Focareta and Cole, 2016; Deryckere et al., 2021; Duruz et al., 2023) which demonstrate a lot of specific features in their complex nervous system. Data about presence and distribution of pan-neuronal proneurogenic and neurogenic factors are scarce in case of other Molluscan groups, particularly in gastropods.

Sox genes, characterized by the presence of the high mobility group (HMG) DNA binding domain, constitute a group of transcription factors with pivotal roles in cell specification and tissue differentiation (Pevny and Placzek, 2005). Among the panoply of Sox genes, it is the SoxB representatives that emerge as key players in neuronal development processes. Their early expression during gastrulation contributes substantially to ectodermal patterning and gastrulation movements (Okuda et al., 2010). During neurogenesis of vertebrates, SoxB1 and SoxB2 family genes maintain the accurate balance between cell proliferation and differentiation, acting as a gatekeeper to inhibit premature differentiation (Bylund et al., 2003; Masui et al., 2007). Moreover, SoxB1 genes play a pivotal role in neural subtypes differentiation within the central nervous system, underlining their significance in the intricate process of neural specification (Pevny and Placzek, 2005; Kiefer, 2007; Panayi et al., 2010). SoxB1 expression occurs in neurogenic zones where it maintains the cells’ ability to proliferate and inhibits further differentiation (Bylund et al., 2003; Masui et al., 2007). In its turn, SoxB2 group genes repress SoxB1 activity and allow progenitor cells to differentiate into neurons (Pevny and Placzek, 2005; Kiefer, 2007).

Sox genes have been identified in many invertebrate species (Phochanukul and Russell, 2010). In addition to conservative features, the specific role of SoxB1 and SoxB2 in the larval development and neurogenesis has been mentioned in different groups. In some cnidaria, SoxB genes act as one of the key regulators of larval morphogenesis (Chrysostomou et al., 2022). In the nematode *C. elegans*, SoxB1 and SoxB2 are largely recruited into the mechanism of the larval to adult transition of the nervous system (Vidal et al., 2015). SoxB expression has been also detected in cnidarians (Magie et al., 2005; Shinzato et al., 2008; Richards and Rentzsch, 2014), flatworms (Dong et al., 2014; Monjo and Romero, 2015), acoels (Semmler et al., 2010), annelids (Kerner et al., 2009; Sur et al., 2020), insects (Buescher et al., 2002; Wilson and Dearden, 2008), and bryozoans (Fuchs et al., 2011).

Despite the emerging understanding of Sox gene functions in various organisms, the specifics of their expression and role remain largely unexplored outside of well-studied models such as *Drosophila*, sea urchins and nematodes. In particular, their functions in the most diverse group of Lophotrochozoans - Mollusca - remain obscure. Recent studies on cephalopods have shed light on SoxB-family gene expression and their involvement in the neuronal precursor specification within head ectoderm and developing ganglia (Focareta and Cole, 2016; Deryckere et al., 2021; Duruz et al., 2023). However, cephalopods have a largely modified development without larvae in their life cycle. Information regarding SoxB gene expression in basal molluscan group possessing true larvae as well as its correlation with larval neurogenesis is scarce.

We use larvae of the freshwater gastropod *Lymnaea stagnalis* (*L. stagnalis*) to study the expression patterns of SoxB1 and SoxB2 in the course of development from gastrulation to hatching. In addition, we apply FMRFamide immunostaining to visualize specific neuronal elements. FMRF- amide is a neuropeptide that is expressed in the earliest neurons of *L. stagnalis* (Voronezhskaya and Croll, 2015). FMRFamide immunostaining has been used in gastropods to reveal central ganglia anlagen (Croll and Voronezhskaya, 1996; Voronezhskaya and Elekes, 1996; Nezlin and Voronezhskaya, 2017), as well as, numerous peripheral neurons and the local neural networks (Croll, 2000; Faccioni-Heuser et al., 2004; Voronezhskaya and Croll, 2015). Parallel visualization of FMRFamide-immunoreactive elements with Sox gene expression allows to correlate location of presumptive neurogenic areas with emerging larval nervous system.

Employing modern techniques such as in situ hybridization chain reaction (HCR-ISH), immunohistochemistry (IHC) for SoxB1 and FMRFamide, and proliferation assays, our research provide a comprehensive outlook into the correlation between SoxB1\SoxB2-expressing cells, the formation of central ring ganglia, and presence of active proliferation zones within the larval body. These findings provide crucial insights into the conserved role of SoxB family proteins in neurogenesis across the evolutionary spectrum. Furthermore, they underscore the crucial distinction in the expanded SoxB1 expression that is specific to gastropod mollusks. This study thus contributes to our broader understanding of Sox gene functions in neurogenesis in non-model organisms and highlights the importance of such studies in diverse species.

## 2 Results

The larval development in L. stagnalis takes place inside egg capsules deposited in egg masses by mature snails. This developmental process typically lasts approximately 11 ± 0.5 days at 25°C. Within a single egg mass, larvae undergo synchronous development through distinct stages (Fig 1a), including early cleavage, blastula, gastrula, trochophore, veliger, metamorphic larvae, postmetamorphic adult-like snails, and hatchlings (Meshcheryakov, 1990; Ivashkin et al., 2015).

**Figure 1.**
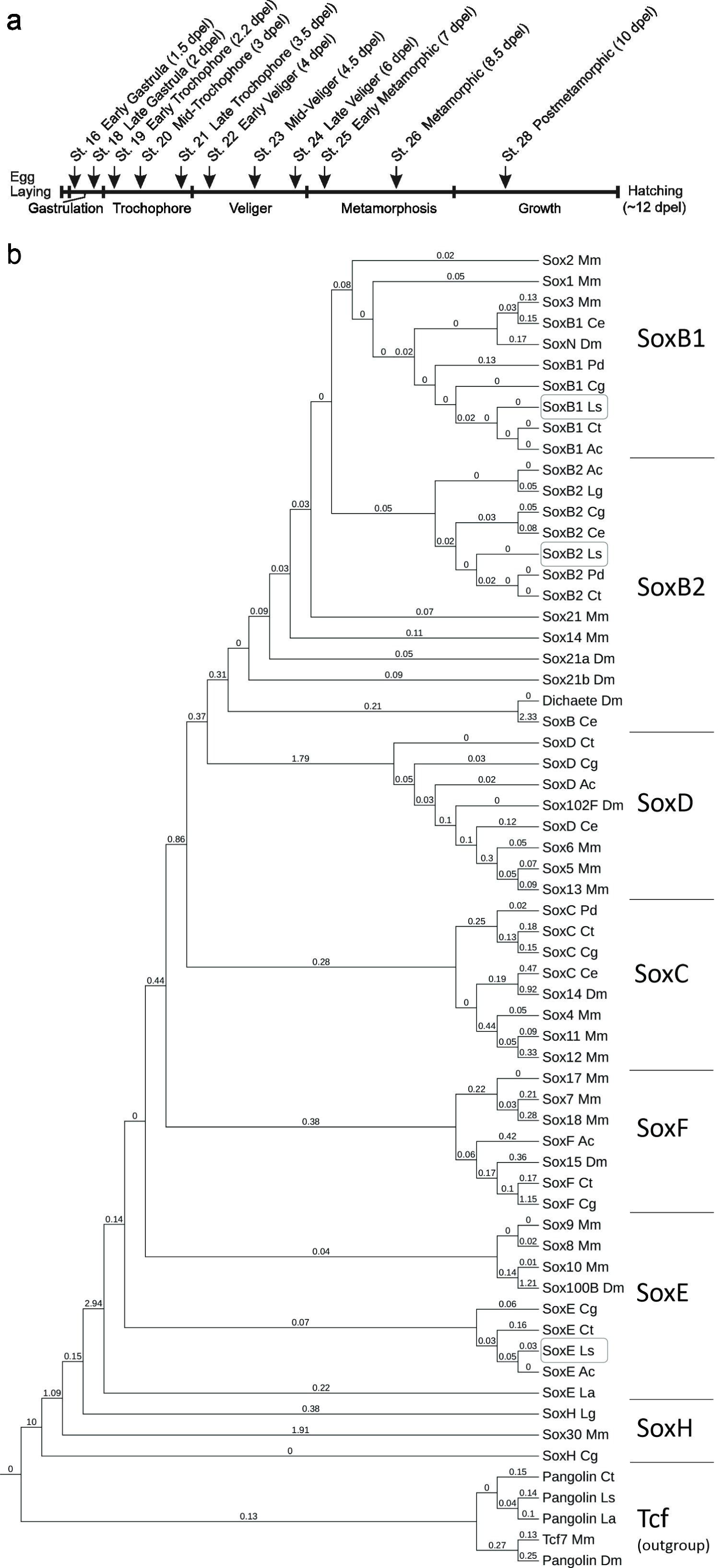
Developmental staging and phylogenetic analysis of the sox gene family in Lymnaea stagnalis. (a) - Developmental periodization of Lymnaea stagnalis and the stages analyzed in the current study. The timing of development is indicated in days post egg laying (dpel). (b) - Maximum likelihood phylogeny of a conserved HMG domains (81 aa) from Sox proteins across various species. Above phylogram braces, the Branch lengths values are written. Detailed species names and sequence accession numbers can be found in Supplementary File, Table S1. The tree utilizes Tcf/Pangolin as an outgroup. The HMG domains of recognized Sox families cluster together in the tree as anticipated and are marked as SoxB1, SoxB2, SoxD, SoxC, SoxF, SoxE, and SoxH. L. stagnalis genes are highlighted within frameworks. Species abbreviations: Ac - Aplysia californica; Ce - Caenorhabditis elegans; Cg - Crassostrea gigas; Ct - Capitella teleta; Dm - Drosophila melanogaster; La - Lingula anatina; Lg - Lottia gigantea; Ls - Lymnaea stagnalis; Mm - Mus musculus; Pd - Platynereis dumerilii.

Our investigation focuses on the stages spanning from the end of gastrulation to hatchlings, specifically honing in on the developmental transition from the trochophore stage, where initial nerve elements emerge, to hatchlings, characterized by the presence of central ring ganglia similar to those in adult L. stagnalis. The central ganglia encompass the esophageal ring ganglia (paired cerebral, pedal, and buccal ganglia), visceral loop ganglia (paired pleural and parietal ganglia, and an unpaired visceral ganglion), and a peripheral osphradial ganglion. It is noteworthy that L. stagnalis possesses an extensive peripheral nervous system, including a plexus in the head, tentacles, and foot. The number of peripheral neurons in this plexus is greater in sum than the number of neurons in the central ganglia (Faccioni-Heuser et al., 2004; Wyeth and Croll, 2011). Gastrulation marks the transition to a pre-neuronal stage characterized by ectoderm patterning. Subsequently, during the trochophore stages, early larval neurons emerge, and ganglia rudiments begin to form. Veliger and metamorphic stages represent the development of the primary central ganglia. Both central and peripheral nervous structures attain mature configurations following metamorphosis and hatching.

### 2.1 Identification of SoxB Genes in the Transcriptome of *L. stagnalis*

To identify SoxB-family genes, we examined the available partial transcriptome of *L. stagnalis*, sourced from a mix of postmetamorphic snails (st. 28-29) and adult nervous systems. Several Sox gene sequences were discerned based on the sequence of their HMG domains. Subsequent phylogenetic analysis classified them into the SoxB1, SoxB2, and SoxE subfamilies, aligning them with their respective orthologs from other Lophotrochozoans. The gene Pangolin, belonging to the Tcf family, was also identified and included as an outgroup. These genes were designated as Ls- SoxB1, Ls-SoxB2, Ls-SoxE, and Ls-Pangolin. The identified SoxB genes distinctly clustered into the SoxB1 and SoxB2 groups (Fig. 1b). Both L. stagnalis SoxB-family sequences contain Sox-family KKDK and LPG conserved motifs. The Ls-SoxB1 sequence has the TKT motif, while the Ls-SoxB2 has the PKS motif within the HMG domain, identifying them as belonging to the SoxB1 and SoxB2 subfamilies, respectively. The obtained sequences were used for the further work.

### 2.2 Specificity of Sox2-like Immunoreactivity for Ls-SoxB1

To conduct a thorough examination of Ls-SoxB1 expression, encompassing potential translational regulation, we employed a dual approach by analyzing both mRNA and protein expression. Fluorescent HCR in situ hybridization (HCR-FISH) for Ls-SoxB1 was combined with immunohistochemistry (IHC) using antibodies targeting mammalian Sox2 (Sox2-IR). To validate the specificity of the antibodies employed, we performed Western blot analysis, confirming the presence of Sox2-IR bands in st. 22 and st 29 L. stagnalis. The observed bands exhibited a molecular weight close to the predicted weight for Ls-SoxB1 based on its sequence (42 KDa; Fig. 2a).

**Figure 2.**
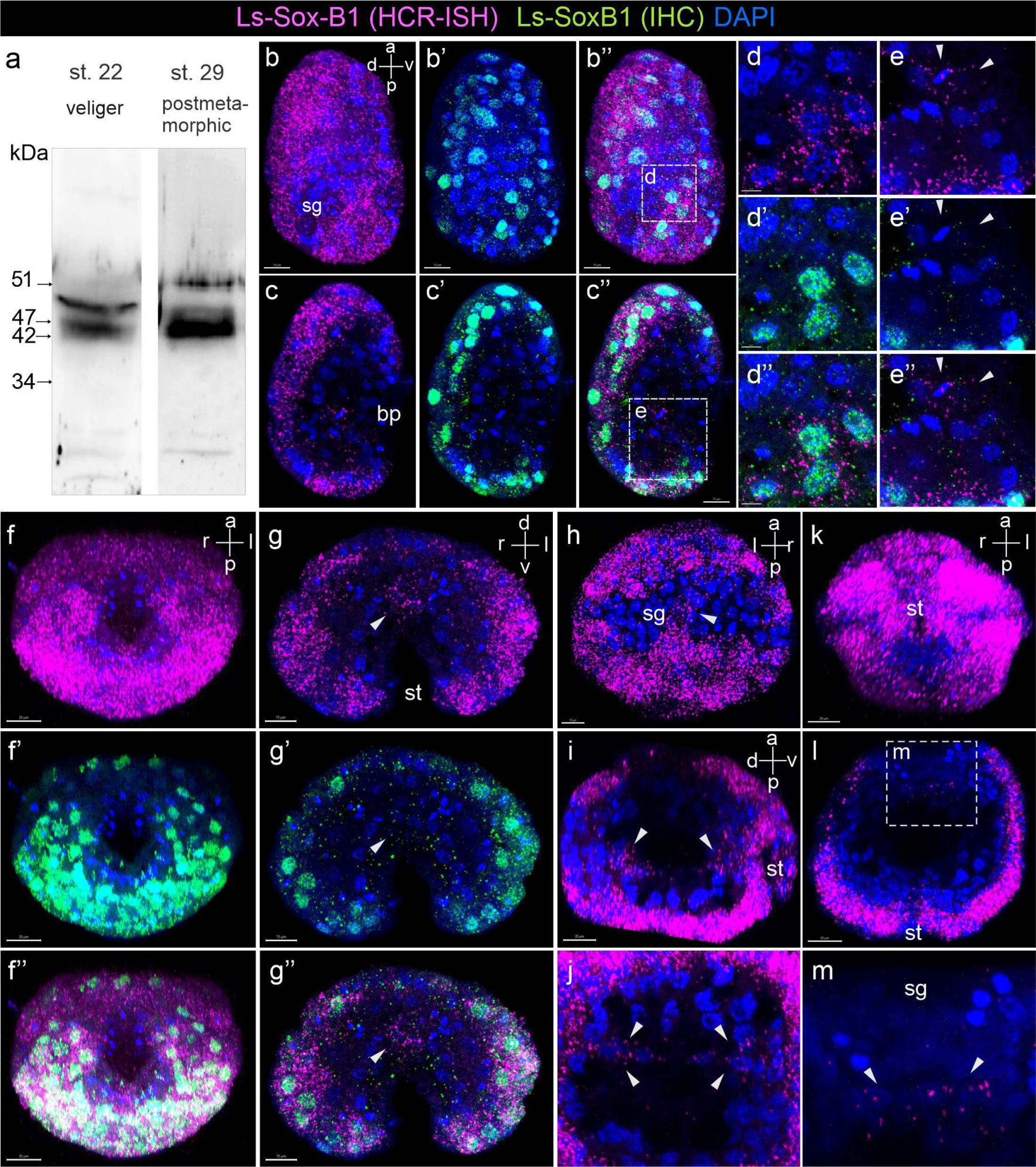
Expression of Ls-SoxB1 in L. stagnalis gastrula and early trochophore larvae. (a) – Western blot using rabbit antibodies against Sox2, showing specific bands in lysates from L. stagnalis veliger larva (st. 22) and postmetamorphic snail (st. 29). (b-b’’) – Maximal projection and (c-c’’) – medial optical sections of the gastrula (st. 17). Both Ls-SoxB1 mRNA and protein are almost ubiquitously distributed in the most ectodermal cells. (d-d”) – Enlarged images demonstrating the cytoplasmic localization of Ls-SoxB1 HCR and the nuclear localization of Ls-SoxB1 IHC signals in ectoderm cells. (e-e”) – Ls-SoxB1 transcripts but not protein are present in the posterior blastopore lip cells (arrowheads). (f-f’’) – late gastrula (st.18), view from the mouth, (g-g’’) – st.19, transverse section through the stomodeum, and (h) view from the forming shell gland of the early trochophore (st.19), arrowheads indicate cells of the gut adjacent to shell gland anlage (i) – sagittal section through the archenteron and (j) – transversal sections through the posterior portion of archenteron in st.19. Arrowheads indicate the Ls-SoxB1 positive streak of small cells along the archenteron wall (arrowheads). (k) – view from the mouth of the st.19. All ectodermal cells maintain a high expression level of Ls-SoxB1, with the exceptions of the zone ventrally to the stomodeum. (l, m) – transverse section of st.19. Ls-SoxB1 expression in the cells of forming gut (arrowheads) adjacent to the shell gland anlage. Abbreviations: bp – blastopore, sg – shell gland anlage, st – stomodeum (mouth).

Subsequently, in situ staining revealed a high concordance between Ls-SoxB1 mRNA expression and Sox2 immunoreactivity (Sox2-IR) in the developing larvae at various stages. Additionally, we identified some specific regions that lacked Sox2-IR but exhibited exclusive Ls-SoxB1 mRNA expression. This observation was attributed to translational regulation, a phenomenon known for SoxB1 orthologs in other animals (Angerer et al., 2005). The specificity of Sox2-IR to SoxB1 was also supported by co-staining with Ls-SoxB2, revealing a distinct expression pattern.

### 2.3 Ls-SoxB1 Expression Prior to Ganglia Formation

During gastrulation and early organogenesis (st. 16-17), particularly at stages encompassing gut formation, Ls-SoxB1 expression predominated in ectodermal regions (Fig. 2b-c’’). Notably, all surface areas exhibiting Ls-SoxB1 expression demonstrated co-occurrence with Sox2-IR, confirming the presence of both mRNA and protein (Fig. 2d-d’’). In addition to ectodermal regions, Ls-SoxB1 expression was observed in the epithelial cells of the posterior blastopore lip, while the symmetrical anterior portion lacked this expression in st. 17 (Fig. 2c-c’’). It is worth noting that while SoxB1 mRNA was present in these specific Ls-SoxB1 positive cells in the archenteron (future foregut), the respective protein was not (Fig. 2e-e’’). At the subsequent stages (st. 18-19), positive Ls-SoxB1 and Sox2-IR persisted in ectodermal cells (Fig. 2f-f’’), with the exceptions of the shell gland formation zone at the dorsal embryo side (Fig. 2h) and the ventral zone around the stomodeum (mouth opening) (Fig. 2f-f’’, k). Ls-SoxB1 expression was observed in a streak of small cells in the ventral part of the forming intestine (arrowheads in Fig. 2i, j), and concentrated in the wall of the forming gut adjacent to the shell gland formation prospective zone in st. 19 (arrowheads in Fig. 2g-g’’, l, m).

At the trochophore stage (st. 20-21), both Ls-SoxB1 expression and Sox2-IR were present in most ectodermal cells, including cephalic plate areas, apical ciliated cells, head vesicles, the entire surface of the foot rudiment, and around the shell gland anlagen (Fig. 3a-d). A continuous streak of small cells expressing Ls-SoxB1 was located on the ventral side along the forming intestine in st. 20 (Fig. 3e). In the early veliger (st. 22) stage, Ls-SoxB1 expression and Sox2-IR decreased in head vesicle cells, velum, and the area forming a transverse ventral groove (Fig. 3f, g, g’), while becoming prominent in the stomodeum (Fig. 3f’). Notably, the cells ventral to the mouth opening, medial ciliated cells in the foot (Fig. 3 h-h’’), as well as the shell gland (Fig. 3 k-k’’), lacked Ls-SoxB1 expression. Expression remained in the ventral portion of the intestine (Fig 3 i-i’’) and notably in the gut adjacent to the shell gland.

**Figure 3.**
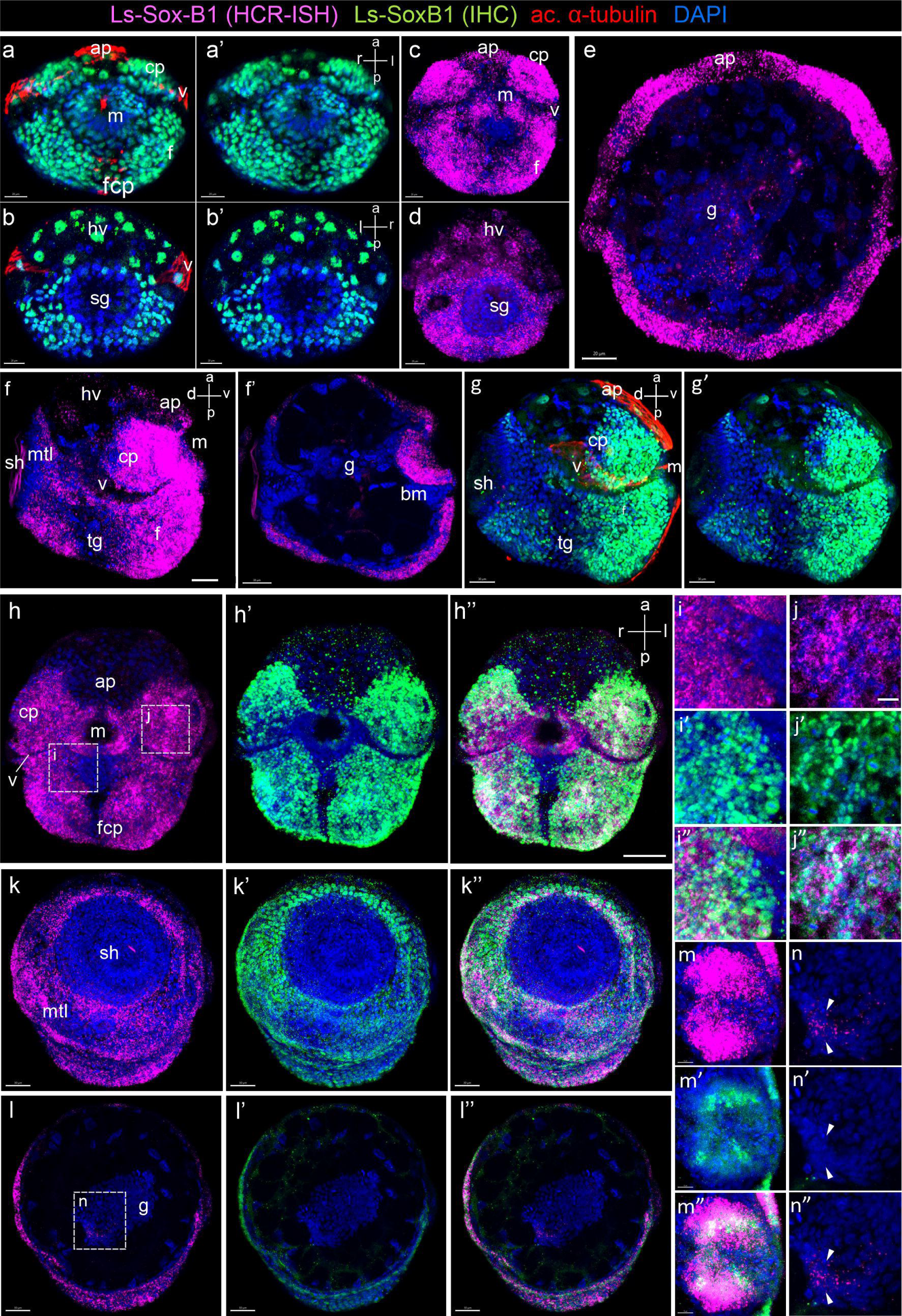
Ls-SoxB1 Expression in L. stagnalis late trochophore and early veliger. (a, a’, c) – oral and (b, b’, d) – shell gland views of late trochophore (st. 20). Ls-SoxB1 transcript and protein observed in cephalic plates, apical ciliated cells, head vesicles, and entire surface ectoderm of foot rudiment. (e) – transverse section of st. 20 showing continuous Ls-SoxB1 plate along the gut. (f, g, g’) – side view of early veliger (st. 22) with sustained Ls-SoxB1 expression, (f’) – sagittal section with prominent expression in oral cavity cells. (h-h’’) – views from mouth, (k-k”) shell gland, and (l- l”) transverse section of st. 22. Note sustained Ls-SoxB1 expression in cerebral plates, foot, and mantle, but not in shell gland. (i-i’’) – high magnification showing decreased Ls-SoxB1 expression and maintaining Sox2-IR in the foot area ventrally to the mouth. (j-j’’) – high magnification showing coexistence of cytoplasmic Ls-SoxB1 and nuclear Sox2-IR within cells of cerebral plate. (m-m’’) – high magnification of oral cavity wall with broader Ls-SoxB1 expression than Sox2-IR zone. (n-n’’) – solitary Ls-SoxB1 expression in the gut cells. Abbreviations: ap – apical plate; bm – buccal mass; cp – cerebral plate; f – foot; fcp – foot ciliary plate; g – gut; hv – head vesicle; m – mouth; mt – mantle; tg – transverse foot groove; sh – shell gland; v – velum.

Detailed examination revealed variations between Ls-SoxB1 expression and Sox2-IR in different larval tissues in st. 22. In particular, there was a gradual decrease in Ls-SoxB1 expression in the foot region ventral to the mouth, accompanied by bright immunoreactivity in the corresponding area (Fig.3i-i’’). In the cells of the cerebral plate, the cytoplasmic expression of Ls-SoxB1 completely coincided with the nuclear localization of Sox2-IR (Fig. 3j-j’’). The zone of Ls-SoxB1 expression in the oral cavity wall exceeded that of Sox2-IR (Fig. 3m-m’’), and a gradual decrease in protein expression was observed from the surface to the depth, extending from the mouth opening to the depth of the intestine in st. 22. Finally, there was exclusive Ls-SoxB1 expression in the gut cells without Sox2-IR (Fig.3 n-n’’). Overall, these results indicate a high plasticity of Ls-SoxB1 product expression at both transcriptional and translational levels.

### 2.4 Ls-SoxB1 Expression During Ganglia Formation and Metamorphosis

Throughout the veliger stage (st. 23), Ls-SoxB1 expression persisted across the extensive ectodermal area of the head, including the tentacles, as well as the dorsal, lateral, and ventral surfaces of the foot ectoderm. Additionally, Ls-SoxB1 expression was observed in the mantle and visceral ectoderm areas (Fig. 4a, f-g’). Notably, Ls-SoxB1 was present in the oral cavity and the dorsal portion of the developing midgut in st. 23 (Fig. 4b, b inset, c), with a few cells located in the region of the cerebral ganglia rudiment (Fig. 4b) and cells in the subepithelial layer in the foot (Fig. 4e). In all these regions, except the midgut, the expression pattern of Ls-SoxB1 coincided with Sox2-IR in st. 23 (Fig. 4a, f-g’). The shell area, ciliated structures (e.g., apical plate, velum, foot medial cells), and cells in the transverse foot groove remained free of SoxB1 expression.

**Figure 4.**
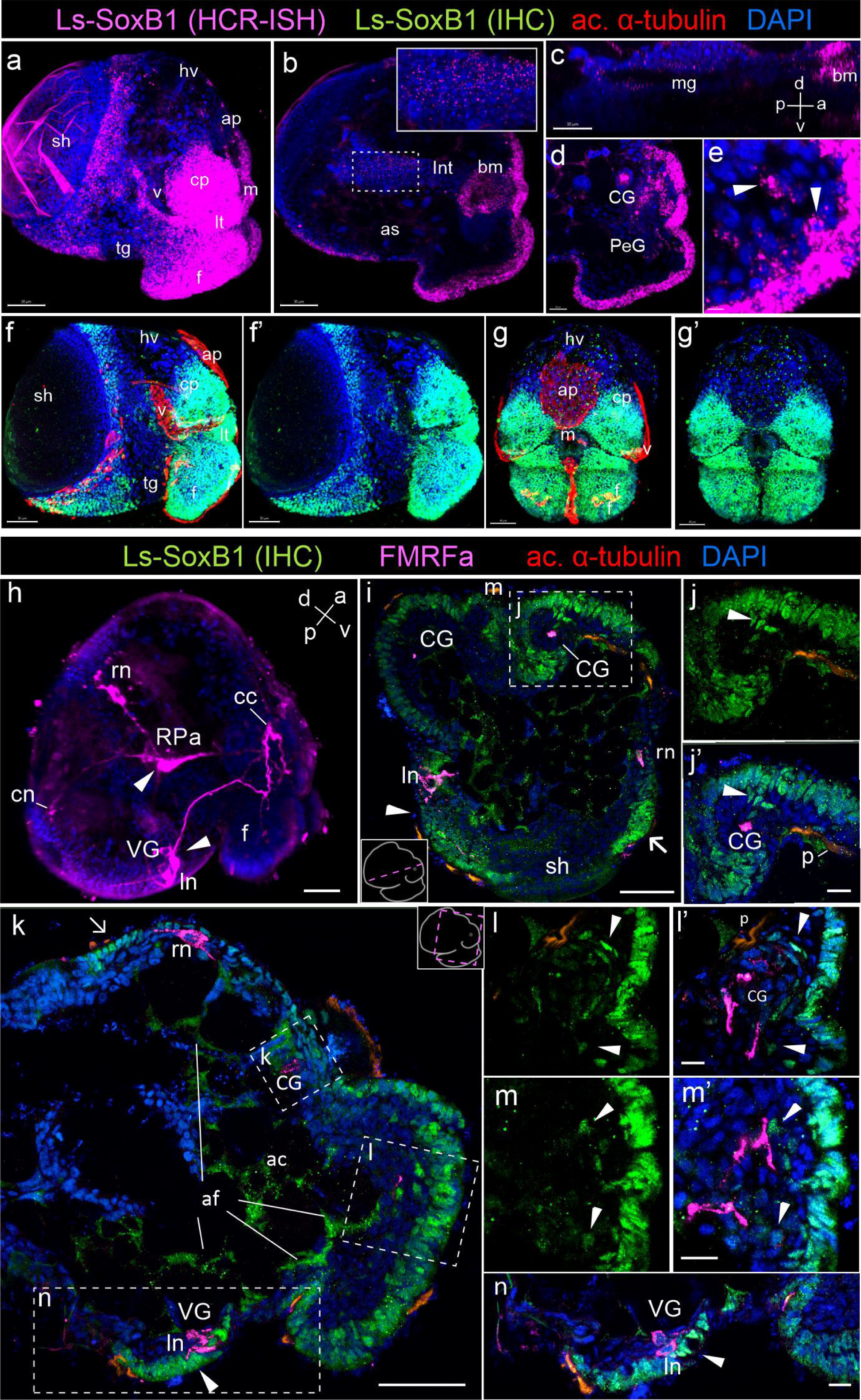
Ls-SoxB1 expression in *L. stagnalis* veliger during emergence of early nerve elements. (a-e) – Ls-SoxB1 expression and (f-g’) – Ls-SoxB1 immunoreactivity in the head and foot ectoderm, in the mantle and visceral ectoderm of veliger (st. 23). Low expression occurs in ciliated apical plate, foot ciliary plate, and foot transverse grove. (b, c) – sagittal section, Ls-SoxB1 expression in stomodeum and forming midgut in st. 23. (d, e) – sagittal sections, Ls-SoxB1 positive cells in the rudiment of the cerebral ganglia and in the subepithelial cells in the foot in st. 23. (h) – whole mount of veliger (st. 23) with early FMRFamide positive peripheral cells, neuropil of ganglia anlagen, and solitary cells within forming parietal and visceral ganglia. (i-n) – sections of st. 23. (i-k) – sections, thick Ls-SoxB1 positive epithelial layer over the forming cerebral ganglion and (n) adjacent to the early FMRF-IR neurons in the visceral part of the body at st. 23. (j-j’) – subepithelial Ls-SoxB1 positive cells beneath the thick Ls-SoxB1 ectoderm of the cephalic plate and (l-m’) in the subepithelial layer in the foot and adjacent to the FMRF-IR nerve fibers at st. 23. Abbreviations: ap – apical plate; af – autofluorescence; as – albumen sac; bm – buccal mass; cc – cerebral commissure; CG – cerebral ganglion; cn – caudal pioneer neuron; cp – cephalic plate; f – foot; hv – head vesicle; int – intestine; ln – left pioneer neuron; lt – labial tentacle; m – mouth; mg – midgut; p – protonephridia; PG – pedal ganglion; rn – right pioneer neuron; RPa – right parietal ganglion; tg – transverse foot groove; sh – shell; v – velum; VG – visceral ganglion.

To further investigate Ls-SoxB1 expression and its association with differentiated neural elements, we conducted combined immunostaining using antibodies against Sox2 and FMRFamide. At the veliger stage (st. 23), FMRFamide-like immunoreactivity (FMRF-IR) highlighted early peripheral cells (caudal, left, and right pioneer neurons), neuropil in ganglia rudiments, and solitary cells within the developing parietal and visceral ganglia (Fig. 4h). Sox2-IR was observed in the thick visceral ectoderm adjacent to the left and right early FMRF-IR neurons (Fig. 4k, n), as well as in the epithelial layer cells above the forming cerebral ganglion in st. 23 (Fig. 4i, k, j, j’). Notably, certain SoxB2-IR cells were located subepithelially between the thick SoxB1-positive ectoderm of the cephalic plate and the cerebral ganglia rudiment (Fig. 4j-j’, l-l’), as well as beneath the SoxB1- positive foot ectoderm in st. 23 (Fig. 4k, m-m’).

At late veliger (st. 24) and metamorphic stages (st 25-26) Ls-SoxB1 expression and Sox2-IR persisted in the outermost ectoderm of the head, labial tentacles, and foot (Fig. 5a-a’’, c-c’, e) and the margin of the mantle (indicated by an arrowhead in Fig. 5c-c’); and appeared in cells of central ganglia rudiments in st. 24 (Fig. 5b). Sox2-IR was detected in the epithelium and some subepithelial cells within the head and foot (Fig. 5d). The presence of all ganglia was distinctly marked by the occurrence of FMRF-IR cells and neuropil in st. 25 (Fig. 5f). Sox2-IR was presented along the periphery of the cerebral and pedal ganglia, in the rudiment of the buccal ganglia (Figs. 5g), and along the cerebro-buccal commissure in st. 25 (arrowhead in Fig. 5g). Sox2-IR cells constitute a continuous layer in the foot ectoderm. In addition, the solitary Sox2-IR cells and cell butches located beneath the ectoderm at a distance of 1-2 cell bodies (Fig. 5g, h). A subset of deeply immersed subepithelial Sox2-IR cells located close to the ganglia was associated with FMRF-IR processes originating from the pedal ganglia in st. 25 (arrowheads in Fig. 5h). Similar epithelial, subepithelial and deeply immersed subepithelial subpopulations of Sox2-IR cells can be selected in the head as well (Fig 5j) while Sox2-IR cells subpopulation separation to distinct layers are most prominent in the foot. Note that FMRF-IR foot sensory neurons did not exhibit Sox2-IR in st. 25 (Fig. 5h). Despite the clear presence of Sox2-IR cells in the cerebral, pedal, and buccal ganglia (i.e., the esophageal ring ganglia), no Sox2-IR cells were detected in the visceral and parietal ganglia in st. 25-26 (visceral loop ganglia; indicated by arrowheads in Fig. 5i, j).

**Figure 5.**
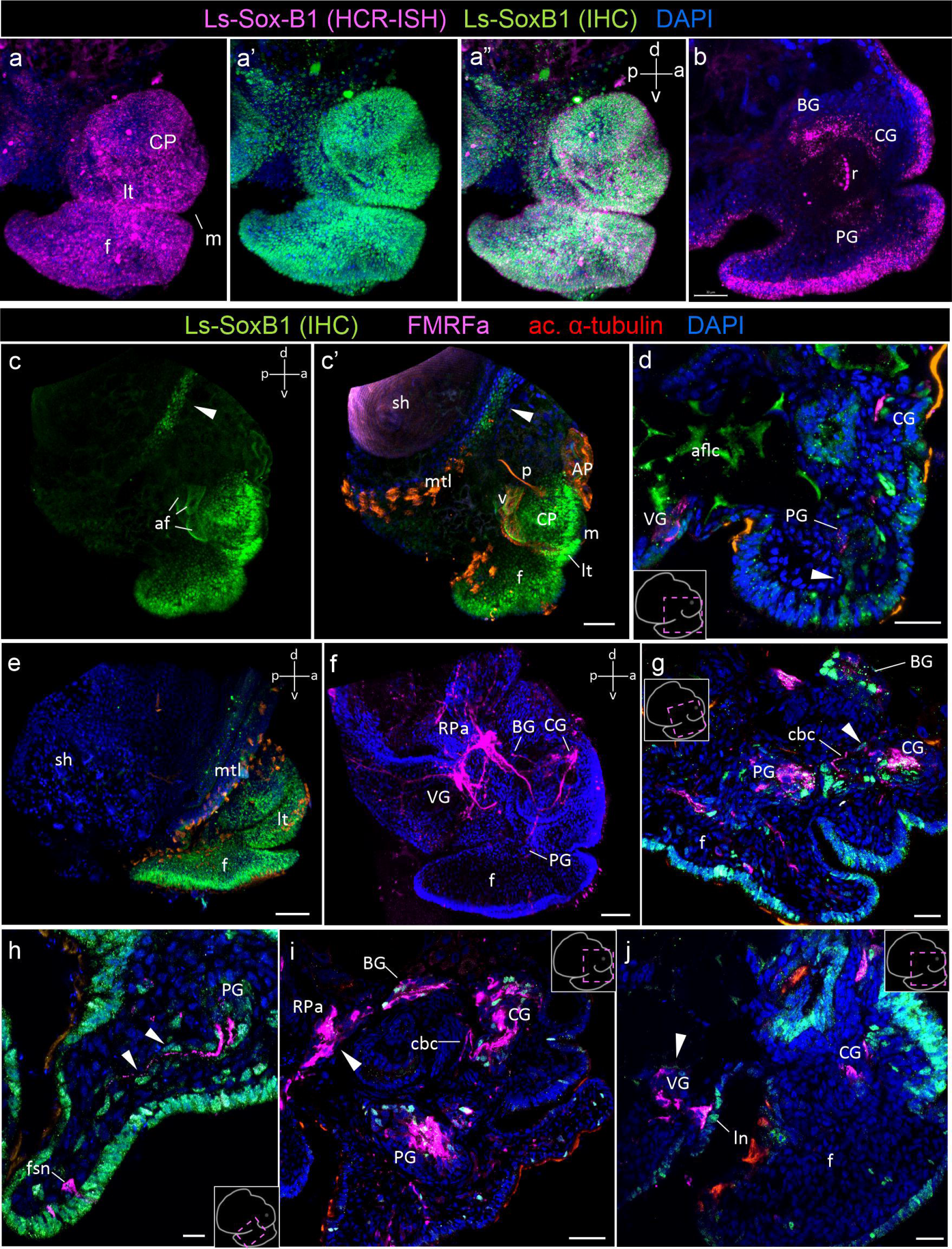
Ls-SoxB1 expression in *L. stagnalis* late veliger and metamorphic larvae. (a-a’’) – whole mount preparation of late veliger (st. 24). Extensive surface distribution of both Ls- SoxB1 transcripts and protein. (b) – parasagittal optical section of st. 24, Ls-SoxB1 expression in solitary cells of cerebral, pedal, and buccal ganglia rudiments in st. 24. (c-c’, e) – whole mount preparation of st. 24. Extensive surface distribution of Ls-SoxB1. Note Ls-SoxB1 positive area at the mantle margin in st. 24 (arrowheads). (d) – cryostat section of st. 24, Sox2-IR in epithelial cells and some subepithelial cells within the foot (arrowheads). (f) – whole mount preparation of metamorphic larvae in st. 25, FMRF-IR marks the main ganglia rudiments and interconnecting connectives and commissures. (g-j) – cryostat sections of st. 25. (g) – Sox2-IR cells in the cerebral, pedal ganglia, and buccal ganglia rudiments, in foot epithelium, in subepithelial layer right underneath the epithelium and deep in the foot in st. 25. (h) – Two subpopulations of Sox2-IR subepithelial cells: right underneath the epithelium and deep in the foot, associated with FMRF-IR fibers (arrowheads) in st. 25 Note Sox2-IR negative FMRF-IR sensory neuron in foot. (i, j) – Sox2-IR cells in the cortical layer of cerebral and pedal ganglia and in buccal ganglia of st. 25 larvae. Note absence of Sox2-IR cells in parietal and visceral ganglia. Abbreviations: AP – apical plate; af – autofluorescence; BG – buccal ganglion; cbс – cerebro-buccal commissure; CG – cerebral ganglion; CP – cephalic plate; f – foot; fns - foot sensory neuron; ln – left pioneer neuron; lt – labial tentacle; m – mouth; mtl – mantle; p – protonephridia; PG – pedal ganglion; RPa – right parietal ganglion; sh – shell; v – velum; VG – visceral ganglion.

At the postmetamorphic adult-like stages (st. 28-29), characterized by fully formed central ring ganglia (Fig. 6d), Ls-SoxB1 expression extends across areas of the head and foot epidermis, including the labial tentacles and the ventral surface of the foot (Fig. 6a, a’, c). In the visceral body region, Ls-SoxB1 expression, in general, decreases at the mRNA level by the postmetamorphic stage, remaining evenly distributed between adjacent cells (Fig. 6b, b’). The epithelium along the edge of the foot sole has a higher level of Ls-SoxB1 expression than in the central zone (Fig. 6c). Numerous neurons at the cortical layer of cerebral and pedal ganglia exhibit Sox2-IR in st. 28 (Fig. 6e, f).

**Figure 6.**
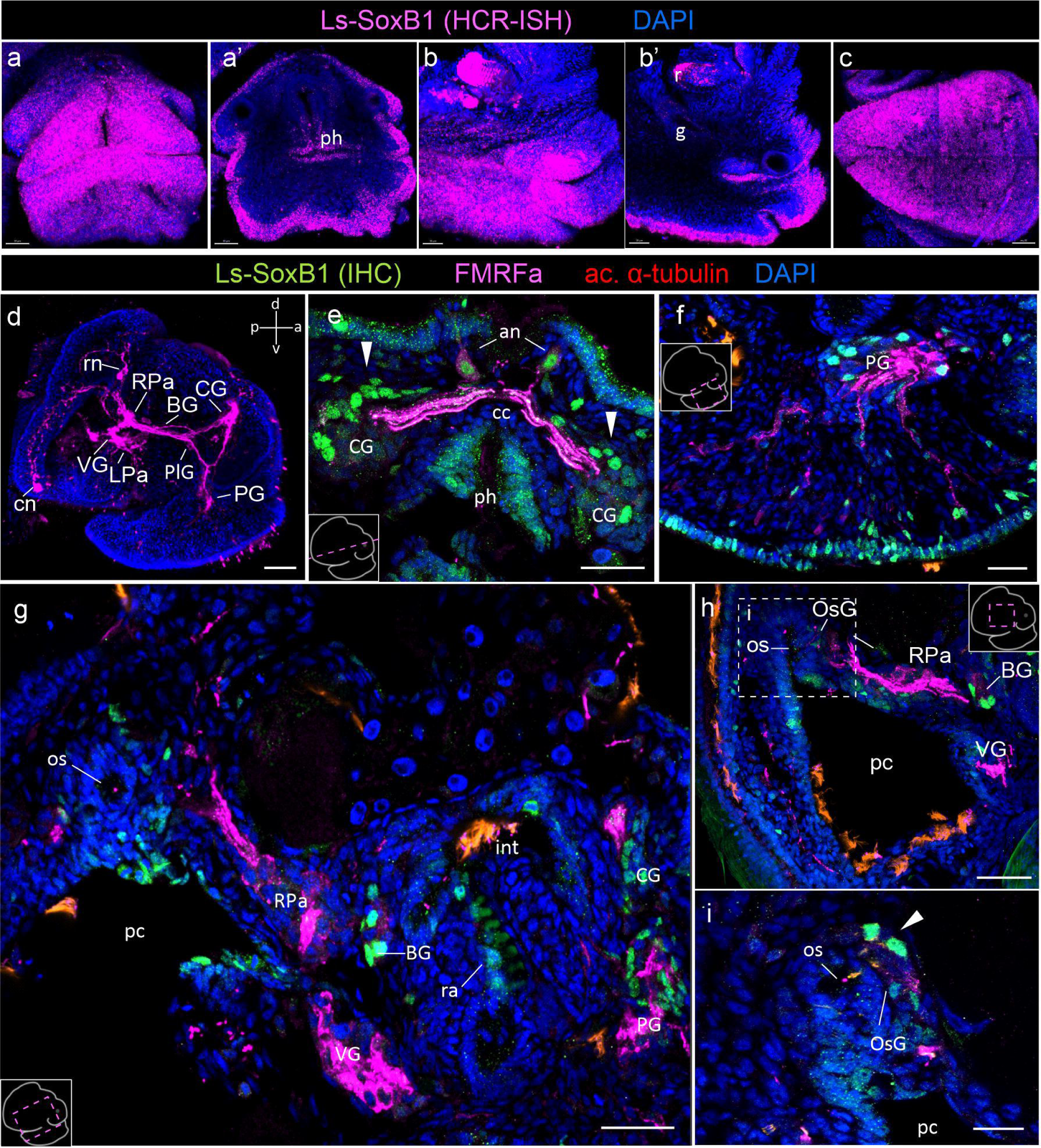
Ls-SoxB1 expression in *L. stagnalis* after metamorphosis. (a, b, c’) – whole mount preparations, (a’, b’) – transverse and sagittal sections of postmetamorphic adult-like stage (st. 28). (a-c) – Ls-SoxB1 maintained in head and foot ectoderm, labial tentacles, and ventral foot surface. Decreased expression in visceral body region and subepithelial layer. (d) – whole mount showing FMRF-IR in central ring ganglia in st. 28. (e) – dorsal view of st. 28 cerebral ganglia with numerous Ls-SoxB1 cells in cortical layer (arrowheads). Note presence of Ls-SoxB1 in apical neurons. (f) – sagittal section through medial part of pedal ganglion in st. 28 with Ls-SoxB1 cells in cortical layer. Note differences in Sox2-IR level in foot epithelial cells. Bright subepithelial Ls-SoxB1-positive cells underneath the epithelial layer and deep in the foot adjacent to FMRF-IR fibers. (g) – sagittal section through the pulmonary cavity with Ls-SoxB1 in cerebral, pedal, and buccal ganglia in st. 28. (h) – high magnification of pulmonary cavity with osphragial ganglion and Ls-SoxB1 cells in st. 28. (i) – high magnification of osphragial ganglion showing two Ls-SoxB1 cells in st. 28. Abbreviations: an – apical neuron; BG – buccal ganglion; cс – cerebral commissure; CG – cerebral ganglion; cn – caudal pioneer neuron; f – foot; g – gut; int – intestine; LPa – left parietal ganglion; OsG – osphradial ganglion; pc – pulmonary cavity; PG – pedal ganglion; PlG – pleural ganglion; ra – radula; rn – right pioneer neuron; RPa – right parietal ganglion; VG – visceral ganglion.

Remarkably, apical sensory neurons display both FMRF-IR and Sox2-IR (Fig. 6e). Sox2-IR in foot epithelial cells demonstrates noticeable differences in the level of expression between adjacent cells in st. 28. Also, Sox2-IR is observed only in a few subepithelial cells. The extensive reaction remains in the subepithelial cells and cell batches of the layer beneath the epithelium. To the contrary, solitary subepithelial Sox2-IR cells associated with pedal FMRF-IR nerve bundles demonstrate lower expression level (Fig. 6f). As at the previous stage, Sox2-IR cell bodies are present in the cerebral, pedal, and buccal ganglia but are notably absent in the visceral and right parietal ganglia in st. 28 (Fig. 6g). Additionally, two Sox2-IR cells appear in the osphradial ganglion (Fig. 6h, i).

### 2.5 Proliferative Activity of Ls-SoxB1-Positive Cells

Given the established role of SoxB1 in sustaining the proliferative activity of proneural cells, it is noteworthy that SoxB1 is a distinctive marker of pre-mitotic neuroblasts (Karnavas et al., 2013; Deryckere et al., 2021), thereby identifying both the primarily proliferative zones in the neurogenic epithelium and secondary proliferative zones in CNS. To assess the proliferative capacity of Ls- SoxB1-expressing cells situated in both the ectoderm and the subepithelial layer, we investigated their ability to incorporate 5-ethynyl-2-deoxyuridine (EdU) and undergo proliferation, as evidenced by immunostaining with antibodies against phosphorylated histone H3 (pH3). This investigation encompassed the late trochophore (st. 21), veliger (st. 23), and metamorphic (st. 25) stages.

Following a 5-minute incubation with EdU, numerous EdU-positive cells were distributed throughout the embryonic body at all examined stages. Notably, at the st. 21 and st. 25, the labeled cells were primarily located within the ectoderm (Fig. 7, a-a’, c-c’), whereas at the st. 23, numerous EdU-positive cells were also found in the subepithelial layer (Fig. 7, b-b’). Importantly, the EdU-incorporating cells were evenly distributed without the formation of distinct zones or clusters of EdU-positive cells in the embryonic body. This uniform pattern was consistent across all developmental stages examined (Fig. 7 a-c’). It’s worth noting that numerous EdU-positive cells were found within the areas of forming ganglia.

**Figure 7.**
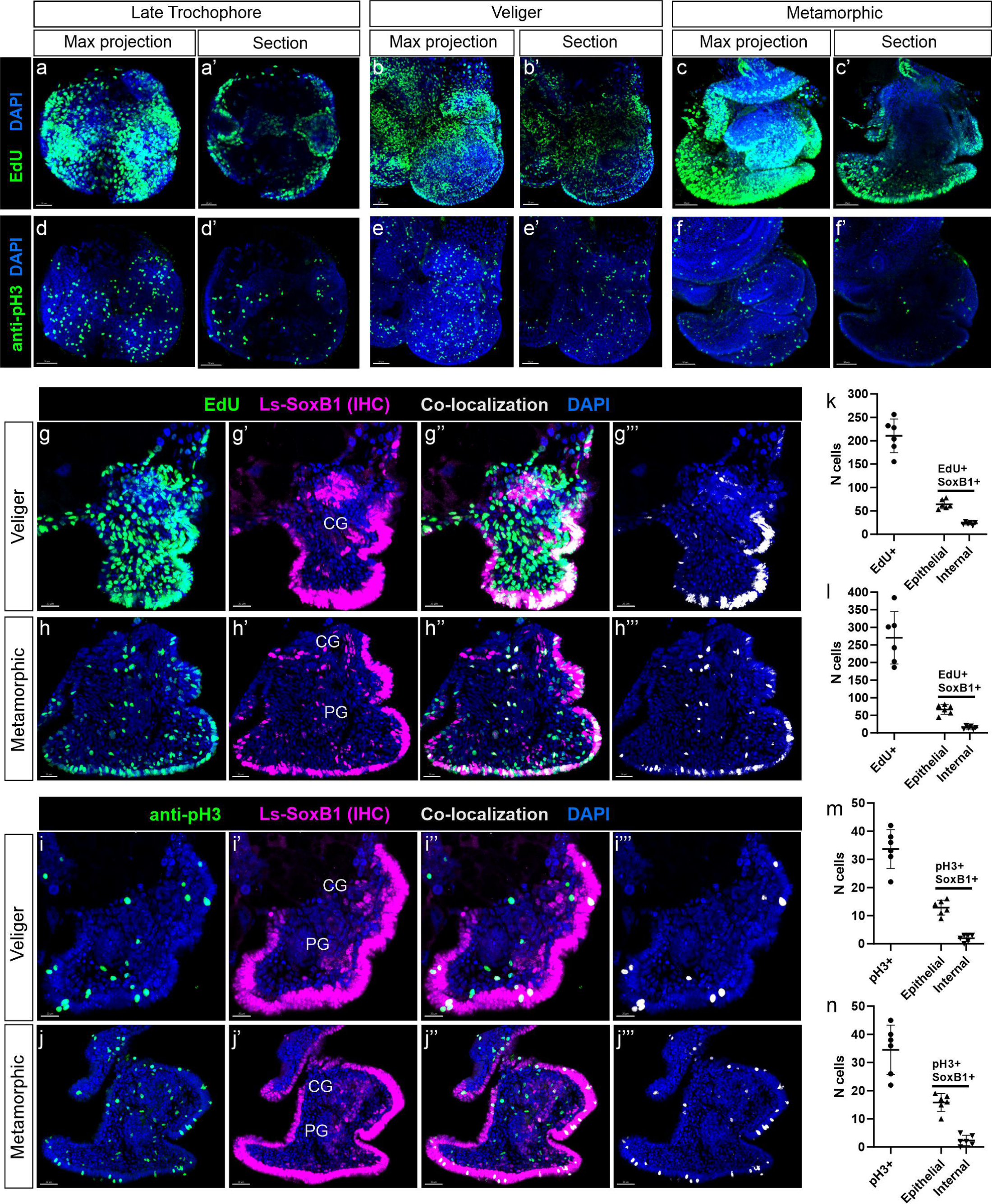
Analysis of Ls-SoxB1 cells proliferative capacity in *L. stagnalis* veliger and metamorphic larvae. (a, b, c) – whole mount and (a’, b’, c’) – sagittal sections of EdU incorporation at st. 21, st. 23 and st. 25, respectively. Numerous EdU-positive nuclei distributed over ectoderm, and occurring in mesenchyme at st. 23. (d, e, f) – whole mount and (d’, e’, f’) – sagittal sections of phosphorylated H3 (pH3) at the same subsequent developmental stages. Positive pH3 nuclei distributed over larval body in st. 21, st. 23 and st. 25. (g-j’’) – Samples of Ls-SoxB1/EdU-positive and Ls-SoxB1/pH3-positive cells with colocalization mainly in epithelium in st. 23 and st. 25. (k-n) – Number of EdU- incorporating, pH3-positive, and Ls-SoxB1/EdU-positive, Ls-SoxB1\pH3-positive cells on sagittal sections at st 23 and st. 25. EdU-incorporating cells prevail pH3-positive cells at both stages. Cells with proliferative activity located mainly in epithelial layer and distributed uniformly at both st. 23 and st. 25. Abbreviations: CG – cerebral ganglion; PG – pedal ganglion.

Further analysis involving double labeling with antibodies against Sox2 and EdU incorporation (Fig. 7 g’-g’’, h-h’’) showed that the majority of epithelial Sox2-IR cells were also EdU-positive, with only a few internal Sox2-IR cells incorporating EdU (Fig. 7 k, l). In both st. 23 and st. 25, some internal Sox2-IR\EdU-positive cells corresponded to the places of forming ganglia.

The immunohistochemical visualization of mitotic cells with antibodies against pH3 revealed a significantly lower number of cells than those incorporating EdU (Fig. 7 d-f’). Similar to EdU labeling, pH3-positive cells exhibited a uniform distribution pattern across the ectoderm and subepithelial layers in st. 23 and st. 25. Importantly, no specific zones with concentrated pH3- positive cells were found at any of the examined stages (Fig. 7 d-f’).

Double labeling with antibodies against Sox2 and pH3 revealed the presence of rare Sox2-IR/pH3-IR cells in both the epithelium and the internal layer (Fig. 7 i-I’’, j-j’’), with a higher prevalence of these cells in the epithelium in st. 23 and st. 25 (Fig. 7 m, n), mirroring the distribution pattern observed for Sox2-IR/EdU-positive cells. Few cells co-expressing Sox2-IR and pH3 were found in a cortical layer of cerebral and pedal ganglia in both st. 23 and st 25.

### 2.6 Ls-SoxB2 Expression and Its Relationship to Ls-SoxB1 Expression

SoxB2, a member of the Sox gene family recognized for its conserved role in establishing neural fate, demonstrates its earliest expression during the late trochophore stage (st 21). By the veliger stage (st 22), the expression pattern of Ls-SoxB2 closely mirrors that of the Ls-SoxB1 gene. Both genes exhibit extensive ectodermal expression, covering broad areas of the head, foot, visceral mass, and mantle (Fig. 8a). Beyond the ectoderm, Ls-SoxB2 is expressed in the ventral part of the oral cavity and the dorsal wall of the radular sac rudiment in st. 22 (Fig. 8a’).

**Figure 8.**
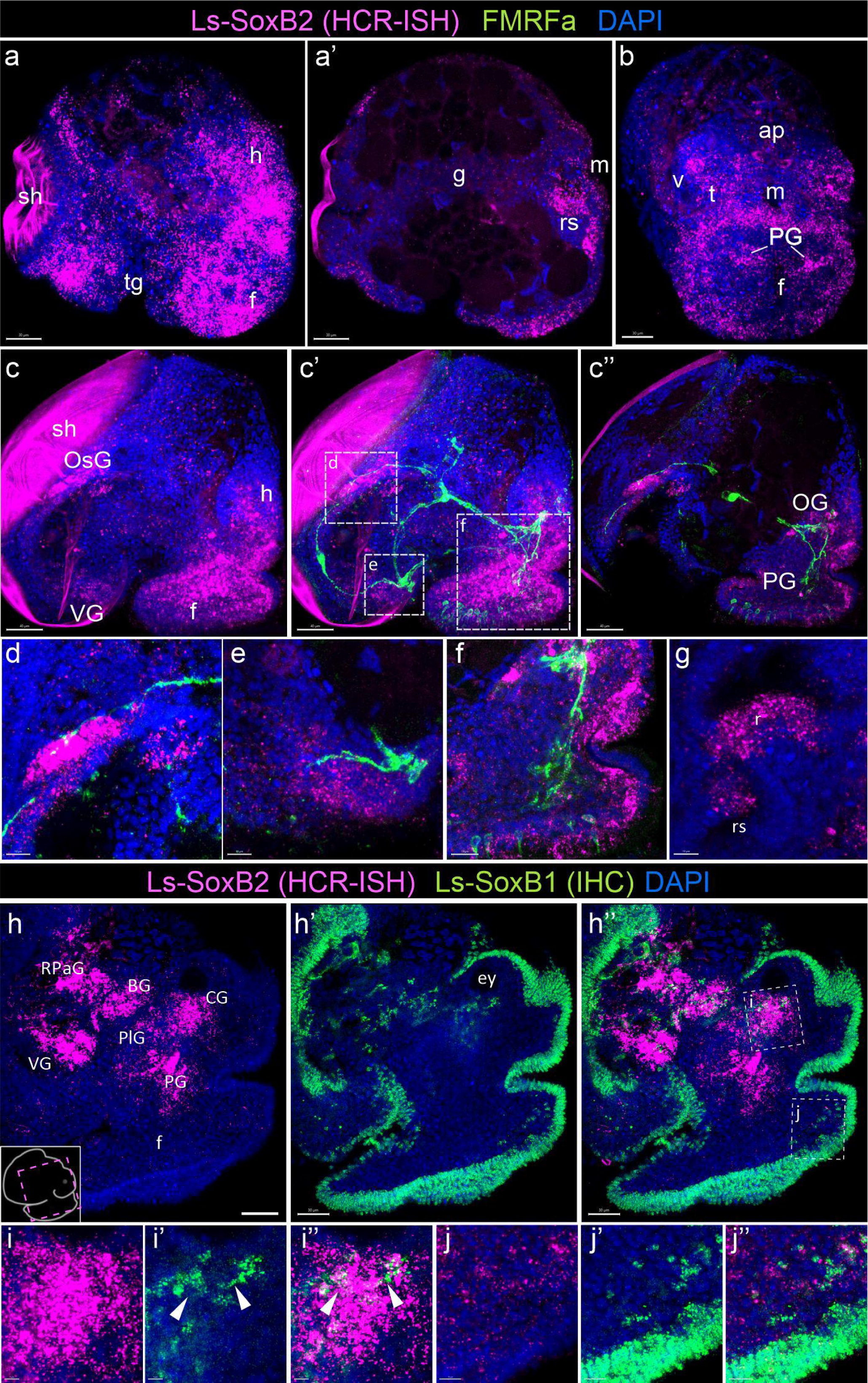
Mutual arrangement of Ls-SoxB2 areas in relation to FMRFamide-containing nerve elements and Ls-SoxB1 expression. (a) – whole mount and (a’) – sagittal section of early veliger (st. 22) showing extensive Ls-SoxB2 expression in head, foot, visceral mass, and mantle ectoderm. (b) – whole mount of st. 23, frontal view with decreased Ls-SoxB2 expression in ectoderm, occurring in pedal ganglia rudiments and subepithelial layer cells of the head and foot. (c, c’) – whole mount and (с”) – sagittal section of st. 25 with limited Ls-SoxB2 expression. Note Ls-SoxB2 positive cells in ganglia rudiments and foot subepithelial layer. (d) - high magnification of osphradial ganglion area. (e) - high magnification of visceral ganglion area. (f) - high magnification of cerebral and pedal ganglia zones. (g) - high magnification of radula region sagittal optical section in st. 25. (h-j’’) - Ls-SoxB1 and Ls-SoxB2 expression at st. 28 larvae. (h-h’) – Ls-SoxB1 preferentially expressed in the epithelium while Ls- SoxB2 concentrated in zones of forming central ring ganglia. (i-i’’) – solitary Ls-SoxB1/Ls-SoxB2- positive cells in cerebral ganglia. (j-j’’) - Ls-SoxB1 and Ls-SoxB2 cells in epithelium and subepithelial layer in the foot. Abbreviations: ap – apical plate; BG – buccal ganglion; ey – eye; CG – cerebral ganglion; h – head; f – foot; g – gut; m – mouth; LPa – left parietal ganglion; OsG – osphradial ganglion; PG – pedal ganglion; PlG – pleural ganglion; r – radula; rs – radular sac; RPa – right parietal ganglion; sh – shell; t – tentacle; v – velum; VG – visceral ganglion.

During the middle veliger stage (st 23), ectodermal expression weakens in ectodermal cells but becomes apparent in subepithelial layer cells of the head and foot, as well as in the pedal ganglia rudiments (Fig. 8b). It is noteworthy that the zone of Ls-SoxB2-positive cells in the foot epithelium loses ubiquitous distribution and forms a rim along the outer edge.

During late veliger and metamorphic stages (st 24-25), Ls-SoxB2 expression is limited to specific ectodermal areas, including the right mantle margin (the region where the osphradial ganglion forms), the ventral part of the visceral mass (adjacent to the forming visceral ganglion), the lateral edges of the foot, and the anterior portion of the tentacles (Fig. 8c, c’). Internalized Ls-SoxB2 positive cells are found in areas corresponding to the pedal, cerebral, and osphradial ganglia rudiments, as well as in the subepithelial layer of foot cells in st.25. Numerous subepithelial Ls- SoxB2 positive cells located approximately one-two cell size beneath the epithelium, some of SoxB2-positive cells also express FMRF-IR (Fig. 8c’’, f). Ls-SoxB2 positive cells are adjacent to FMRF-IR fibers, which mark the osphradial ganglion (Fig. 8d) and visceral ganglion (Fig. 8e), as well as the pedal and cerebral ganglia in st. 25 (Fig. 8f). In the oral cavity, Ls-SoxB2 is expressed in the ventral region of the radular sac (Fig. 8g).

During postmetamorphic stages (st 28-29), Ls-SoxB2 expression almost disappears in the ectoderm but becomes pronounced in subepithelial layer cells and ganglia rudiments. This ganglionic localization of Ls-SoxB2 strongly contrasts with the predominantly epithelial localization of Ls- SoxB1 at st. 28 (Fig. 8h-h’’). Solitary cells expressing both Ls-SoxB1 and Ls-SoxB2 can be found in the cerebral ganglia (Fig. 8 i-i’’) and in subepithelial layer cells in the foot (Fig. 8 i”, k-k’’, j-j’’).

## 3 Discussion

Our study represents the first comprehensive report on the expression of SoxB family genes in the course of gastropod mollusk *Lymnaea stagnalis* development. Our data encompass the entire larval development, from gastrulation to the adult-like snail, including the trochophore and veliger stages, and declare an extended pattern of Ls-SoxB1 expression in the ectoderm maintained both spatially and temporally. Expression of Ls-SoxB1 is supported at both the transcriptional and translational levels, even in postmetamorphic stages that possess fully formed central ganglia and peripheral sensory elements. Significantly, a substantial portion of SoxB1-positive ectoderm is not associated with larval neurogenic areas. In addition, certain cells in the foregut and midgut from the gastrula stage onwards express Ls-SoxB1 solely at the mRNA level and do not contain the corresponding protein. During ganglia formation and axonogenesis, only single specific cells within ganglia and the solitary cells in foot express Ls-SoxB1. Meanwhile, the extended expression of Ls-SoxB2 in the ectoderm is rapidly terminated at veliger stage and later remains in differentiating neurons in ganglia anlages and in subectodermal cells of the foot. These results indicate that the expression pattern of the SoxB transcription factor family in the *L. stagnalis* differs significantly from other investigated vertebrates.

### 3.1 Similarities and Differences with Other Animals

In most bilaterian animals, ectodermal SoxB1 expression during organogenesis is linked to the patterning of the neurectoderm and subsequent neurogenesis (Hartenstein and Stollewerk, 2015). In non-bilaterian cnidarians, as well as bilaterian platyhelminths lacking solid neuroectodermal zones, SoxB1-positive cells are scattered throughout the ectoderm without forming a continuous layer (Magie et al., 2005; Semmler et al., 2010; Richards and Rentzsch, 2014; Monjo and Romero, 2015). In annelids and arthropods, SoxB1 expression in the early stages of neurogenesis is limited to the neuroectodermal placodes, which are distinctly separated from the surrounding ectoderm and lie directly above the emerging structures of the central nervous system (Buescher et al., 2002; Simionato et al., 2008; Wilson and Dearden, 2008; Kerner et al., 2009; Sur et al., 2020). In all cases described, the neuroectoderm-committed areas are characterized by the presence of actively dividing cells.

However, pattern of SoxB1 expression in ectoderm of gastropod *L. stagnalis* contrasted with that in annelids and arthropods, in which it concentrated in clearly delineated and limited areas. In *L. stagnalis*, Ls-SoxB1 expression occupies most of the head, foot, and visceral complex ectoderm. Our observations in *L. stagnalis* indicate that Ls-SoxB1 expression definitely covers presumptive neurogenic epithelium zones starting from the late trochophore and early veliger stages. Increased levels of Ls-SoxB1 protein expression occur in the cerebral plates (cerebral ganglia anlagen), the ventral surface of the forming foot (pedal ganglia anlagen), and body wall near the areas of visceral, parietal, and osphradial ganglia anlagen. Moreover, Ls-SoxB1 remains in ectoderm by late postmetamorphic stages when the most larval ganglia and the most sensory periphery is already formed, and thus is not coincide with known events of nervous system differentiation. Such extensive widespread and prolonged expression of Ls-SoxB1 in *L. stagnalis* suggests that in mollusks, SoxB1 may have an additional function beyond its role in neuroectoderm commitment. Such roles of SoxB1 in some non-neurogenic cells are known both for invertebrates and vertebrates. For example, in *Drosophila*, SoxNeuro (ortholog of SoxB1) takes part in shaping the denticles in the embryonic epidermis (Rizzo and Bejsovec, 2017). In *Xenopus*, Sox3 (one of SoxB1 orthologs in vertebrates) is an important factor in the differentiation of non-neural ectodermal cells in the neural plate border (Schock et al., 2023). However, unlike in the gastropod, these events take place in limited sites of the embryonic epithelium only, and don’t expand to the later developmental stages.

Ls-SoxB1 is also present in zones corresponding to differentiating neurons at stages from late veliger to metamorphosis. However, Ls-SoxB1-positive cells never constitute the majority in the ganglia at any stage. Noteworthy, parietal and visceral ganglia contain Ls-SoxB1-positive processes only but not the Ls-SoxB1 neural cell bodies. This observation aligns with studies in cephalopods, where SoxB1 transcripts were highly expressed in cerebral and pedal cord derivatives but absent in the palliovisceral cord (Focareta and Cole, 2016; Deryckere et al., 2021). Altogether, these results indicate that the late role of SoxB1 in the differentiation of neurons in mollusks is restricted to certain neuronal subtypes and is not attributed to all neurons.

In addition to the role in the development of central nervous system, SoxB1 genes are known to be involved in the differentiation of peripheral sensory cells (Ross et al., 2018). Earlier, the broad expression of SoxB1 and SoxB2 in various epithelial zones of the cuttlefish *Sepia officinalis* has been attributed to the extensive development of peripheral sensory structures in cephalopods (Buresi et al., 2014; Focareta and Cole, 2016). We observed extensive Ls-SoxB1 expression in the epithelial areas of gastropod *L. stagnalis* throughout larval development, including the post-metamorphic stages when the animal already has numerous differentiated peripheral sensory cells. Moreover, the expression in these late larval stages was evenly distributed, so that no restriction to differentiating neural elements could be recognized, as is the case in other animals. In the *Drosophila* trunk sensory zones, for example, the expression of SoxNeuro demonstrates a solid full epithelium expression pattern only at the earlier stages of development. As the differentiation proceeds to sensory cells, sensory neurons, satellite cells, and covering epithelial cells, SoxNeuro expression gains a punctate pattern and gets restricted to pro-neurogenic cells only (Rizzo and Bejsovec, 2017). We do not observe a confinement of Ls-SoxB1 expression to differentiating sensory elements in gastropods.

This observation implies that mollusks exhibit distinctive features in the process of peripheral sensory cell differentiation. In the deuterostome sea urchin, SoxB1 plays an important role in neurogenic differentiation (Feuda and Peter, 2022). Its expression covers the entire ectoderm at the time of larval neurogenesis, but is not confined to the proneurogenic zones only. SoxB2, in turn, shows a key function in the neurogenesis of the sea urchin and is restricted to differentiating neuroblasts (Anishchenko et al., 2018). This interaction between SoxB family genes in echinoderms is strikingly similar to the expression patterns of Ls-SoxB1 and Ls-SoxB2 that we observe in gastropod *L. stagnalis*.Furthermore, Ls-SoxB1-positive cells in the invaginating endoderm in specific areas of the foregut and midgut, and cells in the oral cavity wall show expression exclusively at the mRNA level in the larvae of *L. stagnalis*. The same SoxB1 regulation at the translational level has been described in the endoderm of sea urchins (Wei et al., 2011), suggesting deep analogies between sea urchins and gastropod larvae. The origin of such parallelism is unclear and requires further investigation.

SoxB1 genes play a crucial role in maintaining a proliferative neurogenic state, extending beyond the neuroepithelium. Specifically, Sox2 is known for sustaining the stem cell status within proliferative zones in the central nervous system and retina throughout postnatal development and in adult vertebrates (Martínez-Cerdeño and Noctor, 2018). Secondary proliferative activity within the central nervous system is also observed in flatworms and some arthropods (Hartenstein and Stollewerk, 2015). In the nematode *C. elegans*, SoxB1 is not implicated in the differentiation of the majority of neurons and their epithelial progenitors. Instead, SoxB1 is required to maintain the developmental potential of blast cells generated in the embryo. These cells then divide and give rise to some differentiated neuronal cell types only post-embryonically (Holmberg et al., 2008). In gastropod, we observed that EdU incorporation reflecting DNA synthesis happens in SoxB1-positive cells within ganglia. A few of pH3-positive cells in the cortical layer of the cerebral and pedal ganglia also express SoxB1. This finding aligns with the fact that proliferation of proneurogenic cells within the ganglia in pulmonate gastropods described earlier (Anisimov and Kirsanova, 2002). In cephalopods, some divisions of SoxB1-positive cells have been described in cells along the migration pathway from the surface to the ganglia as well as within the ganglia themselves (Deryckere et al., 2021).

Incorporation of EdU and phosphorylation of histone H3 in gastropod neurons are also anticipated to happen due to endomitosis events that accompany the process of neuron hypertrophy. Neurons in ganglia exhibit a gradual accumulation of ploidy throughout their lifespan, starting from early neurogenesis until the end of life, and can reach a ploidy of 16384C (Kirsanova and Anisimov, 2000, 2001; Anisimov, 2005). In *L. stagnalis*, we observed a lower count of SoxB1 and pH3-positive cells compared to SoxB1 and EdU-positive cells in the developing ganglia. Moreover, only a minimal number of cells within the ganglia, whether SoxB1-positive or SoxB1-negative, expressed pH3. These findings suggest that the endomitotic nuclear divisions accompanying neuronal polyploidy in *L. stagnalis* may not be closely associated with extensive histone H3 phosphorylation.

An intriguing result is the uniform distribution of cell divisions in the epithelium of *L. stagnalis*. We did not observe regions with concentrated proliferating cells in presumptive neurogenic zones at any of the analyzed developmental stages. This indicates that, in contrast to annelids, insects, and cephalopods, cell divisions (including presumptive neuronal precursors) in the neurogenic epithelium of gastropods probably occur over an extended period, and thus, cell divisions are spread in time without forming any clear proliferation-reach domain at any stage of development.

### 3.2 SoxB-Family Genes Expression, Early Neurogenesis, and Evolution of Mollusca

Data concerning early neurogenic events in gastropods and Mollusca phylum in general are currently limited. Only a few morphological studies, focusing on nudibranch *Aplysia californica* and *Melibe leonina*, have described the processes of ectodermal cell delamination during the development of the central nervous system (CNS) ganglia (Kandel et al., 1980; Jacob, 1984; Page, 1992). In these works, the authors reported the development of cerebral ganglia from the ectoderm of cephalic plates and visceral, osphradial, and interstitial ganglia from the adjacent zones of visceropallial ectoderm. However, no molecular-neurogenesis studies have been conducted on this subject.

SoxB1 expression at different developmental stages has been described for several mollusks, including gastropods. In trochophore larvae of the marine gastropod *Patella vulgata*, SoxB1-positive cells were located in the ectoderm regions of the head and trunk, which give rise to the structures of the central nervous system and sense organs (Le Gouar et al., 2004). In *Lottia goshmanii*, another representative of the Patellogastropoda, SoxB1 expression has been described in compact zones on the ventral posterior surface of the trochophore (Tan et al., 2022). The role of SoxB1 and SoxB2 in neurogenesis has been most thoroughly studied in cephalopods, *Octopus vulgaris* and *Sepia officinalis* (Buresi et al., 2016; Focareta and Cole, 2016; Deryckere et al., 2021; Duruz et al., 2023). The authors of all these papers mention that SoxB1 exhibits broad ectodermal expression at early neurogenic stages and further in development. SoxB1-positive ectodermal regions in cephalopods are accompanied by epithelial thickenings, which are hypothesized to give rise to neuronal precursors of the centralized brain (Deryckere et al., 2021). Single-cell transcriptomic data suggest the presence of SoxB1 both in neural migrative precursors together with the expression of Ascl and in some maturing neurons (Duruz et al., 2023).

Cephalopods are thought to have originated from a monoplacophoran or gastropod-like ancestor, thus sharing several developmental and anatomical features with gastropods (Shigeno et al., 2010). The similarity of expanded Ls-SoxB1 expression in gastropod *L. stagnalis* to that in cephalopod mollusks is intriguing. One of the well-known conservative features of SoxB1 proteins is their ability to inhibit the activity of pro-neural factors and maintain pre-neural cells as undifferentiated precursors, retaining their ability to produce neuroblasts (Holmberg et al., 2008). Thus, the widespread expression of SoxB1 in cephalopods is usually attributed to their notably higher number of neurons in the brain compared to other lophotrochozoans (with cephalopods having over 500 million and gastropods around 11 thousand neurons in Lymnaea). It is reasonable to assume that numerous neurons that make up the cephalopod brain arise from these expansive neurogenic zones (Deryckere and Seuntjens, 2018). Unexpectedly, we found a similarly extensive expression of Ls-SoxB1 in ectodermal areas in gastropod, although the number of neurons in their ganglia is considerably fewer than in cephalopods. This fact is consistent with the expression of SoxB2, which is mostly restricted to cells with neurogenic fate during neurogenesis. In vertebrates, the SoxB2-family gene Sox21 represses Sox3 (SoxB1 ortholog) and promotes terminal neural differentiation (Karnavas et al., 2013). Therefore, SoxB2 in mollusks may be a part of the mechanism that limits the number of central ganglion neurons generated during larval development. On the other hand, the expanded expression of SoxB1 in gastropod mollusks may constitute a component of a preadaptation complex that, having originated long ago, underlies the emergence of the sophisticated cephalopod brain and other distinctive traits during evolution. Further studies of molecular neurogenesis in the gastropod larval development will shed light on the fascinating question of the evolution of the molluscan nervous system.

Another similarity between gastropods and cephalopods concerns the presence of SoxB2 in skeletogenic structures. In *L. stagnalis*, Ls-SoxB2 expression occurs in the epithelium of the developing radular sack. The presence of SoxB2 has also been documented in the epithelium responsible for the formation of skeletal oral structures in cephalopods (Focareta and Cole, 2016). This observation lends support to the idea of a common origin of the radula of gastropods and the beak of cephalopods. Going further, we can infer more distant parallels to vertebrates, considering that Sox21 (one of the genes of the SoxB2 family) is expressed in the epithelium of developing teeth and plays a crucial role in the development of tooth enamel in mammals (Saito et al., 2020).

We observed an expanded and prolonged epithelial expression of SoxB1 in the larvae of *L. stagnalis*. Notably, the broad SoxB1 expression characteristic of the ectoderm in gastrulating embryos is also maintained in the epidermis of the post-metamorphic adult-like animal. This retention of embryonic features in later developmental stages suggests a phenomenon akin to neoteny, where an organism preserves characteristics of a younger stage as it progresses through development. In this context, neoteny refers to the protracted retention of embryonic traits in post-metamorphic stages, resulting in a slower ontogeny compared to related animals. This concept parallels the idea that neoteny may contribute to enhanced cognitive abilities, as seen in human evolution with a prolonged period of heightened neuronal plasticity (Somel et al., 2009). The observed alteration in SoxB1 expression aligns with the concept of “transcriptional neoteny,” wherein gene expression in the adult organism mirrors that of an earlier developmental stage, accompanied by a shift in the regulation of corresponding developmental processes (Bakken et al., 2016). In various animals, including representatives of lophotrochozoans, the broad ectodermal expression of SoxB1 is typically limited to the pre-gastrulation and gastrulation stages, and becomes confined to neurogenic zones later in development (Okuda et al., 2010). Consequently, our finding that the gastropod retains a broad SoxB1 expression zone at late developmental stages can be considered a form of transcriptional neoteny. This phenomenon contributes to the complex formation of pedomorphosis in gastropods, where numerous developmental traits can be interpreted as morphological expressions of heterochronic processes, with neoteny playing a crucial role in the diversification of the Mollusca clade (Beklemishev, 1958a, 1958b; Lindberg, 1988). It also aligns with the general neoteny hypothesis on the mollusks origin, previously discussed on the basis of purely morphological evidence (Garstang, 1928; Vagvolgyi, 1967).

### 3.3 Conclusion and Future Directions

In conclusion, our study unveils a nuanced and dynamic expression pattern of SoxB-family genes in the gastropod *L. stagnalis*, offering insights into the potential role of SoxB1 and SoxB2 in neurogenesis and morphogenesis in gastropods. The intriguing parallels with cephalopods and the unique expanded and prolonged SoxB1 expression pattern observed in gastropod *L. stagnalis* open up opportunities for further comparative and evolutionary studies to understand the molecular basis of neural development in mollusks. As we delve deeper into the intricacies of SoxB gene expression and its implications, we anticipate that this research will contribute to the broader field of evolutionary developmental biology.

## 4 Materials and Methods

### 4.1 Animal Handling

The freshwater pond snail *Lymnaea stagnalis* (*L. stagnalis*) is a pulmonate gastropod mollusk. The laboratory population of *L. stagnalis* at the Institute of Developmental Biology RAS originated from Vrije Universiteit, Amsterdam, in 1994. Mature snails were maintained under stable conditions (22– 23°C, 16-8 h light-dark cycle) and provided with lettuce ad libitum. Egg masses were collected daily and examined under a dissecting microscope, with stages of embryonic development determined based on a comprehensive set of morphological characteristics following the method established by Meshcheryakov (Meshcheryakov, 1990).

### 4.2 The RNA Extraction, RNA-Seq Library Preparation, and Sequencing Procedures

*L. stagnalis* postmetamorphic snails (st. 28-29) and adult nervous systems were utilized for total RNA isolation using the RNeasy Mini Kit (Qiagen, Hilden, Germany) in accordance with the manufacturer’s instructions.

The quality assessment of total RNA was performed using the Bioanalyzer 2100 (Agilent, Santa Clara, CA, USA). The quantity and purity of RNA were determined on a NanoPhotometer (Implen). For library construction, 500 ng of total RNA with a RIN ≥7 was employed. The NEBNext® Poly(A) mRNA Magnetic Isolation Module and NEBNext® Ultra II™ Directional RNA Library Prep Kit for Illumina (New England Biolabs, Ipswich, MA, USA) were used based on the manufacturer’s instructions.

The quality verification of the libraries was conducted using the Bioanalyzer 2100 (Agilent, Santa Clara, CA, USA), and the yield was validated through qPCR. Subsequently, the libraries were subjected to sequencing on HiSeq2500 (Illumina, San Diego, CA, USA) with pair-end 126bp readings for transcriptome assembly.

### 4.3 *De Novo* Transcriptome Assembly and Analysis

Raw transcriptome assembly was carried out using the Trinity assembler (v 2.5.1) based on 5 paired- end libraries with 126+126 bp reads. The resulting set of transcripts underwent completeness analysis using the Busco software (v 2.0), utilizing the core Metazoa proteins dataset (n=978).

To address the issue of overabundance inherent in raw transcriptome assemblies (characterized by a substantial number of duplicated transcripts), we implemented expression- and length-based filtration. Additionally, only the longest isoform for each Trinity-derived ‘gene’ was retained.

Abundance estimation involved the use of bowtie-2 for mapping and RSEM for calculating expression values. Subsequently, structural and functional annotation of the transcriptome was performed using Transdecoder (v 5.0.2) and Trinotate (v 3.1.1) tools.

### 4.4 Phylogenetic analysis

Sox protein sequences were identified through keyword searches in the NCBI GenBank (https://www.ncbi.nlm.nih.gov/genbank/) (see Supplementary File, Table S1 for sequences accession numbers). Coding sequences for L. stagnalis were gathered by performing BLAST on molluskan and other lophotrochozoan HMG domains of Sox-family sequences against the L. stagnalis partial transcriptome using BLAST+ v. 2.11 software (Camacho et al., 2009). Putative Sox sequences identified via BLAST hits were translated using the Expasy Translate Tool (https://web.expasy.org/translate/). The HMG-containing open reading frames identified in the L. stagnalis coding sequences were then subjected to protein alignment using AliView v. 1.27 (Larsson, 2014) and the MAFFT multiple sequence alignment method (Katoh and Standley, 2013). Sequences missing part of the HMG domain were excluded from the analysis. The resulting alignment was utilized to calculate the phylogeny in IQ-tree v. 1.6.12 (Trifinopoulos et al., 2016), with Tcf-family proteins serving as an outgroup (Focareta and Cole, 2016). The LG+G4 phylogenetic evolution model was determined using IQ-tree protein Model Finder (Kalyaanamoorthy et al., 2017). Phylogenetic trees were constructed using IQ-tree with ultrafast bootstrapping (Nguyen et al., 2015). Tree visualization was performed using iTOL v. 6.8.1 (Ciccarelli et al., 2006).

### 4.5 HCR Fluorescent *in situ* Hybridization

The HCR probe pools for the fluorescent in situ mRNA visualization of Ls-SoxB1 and Ls-SoxB2 were meticulously generated using the modified HCR 3.0 *in situ* probe generator (Kuehn et al., 2022). To ensure optimal performance, the probe design incorporated filtration against stable secondary structures. Probes were synthesized in abundance, and potential off-target hybridization was rigorously screened using BLAST+. DNA pools, sourced from Synbio, Inc. (see Supplementary File, Table S2 for probe sets sequences), were dissolved in Tris-EDTA prepared with DEPC-treated DNase/RNase-Free MilliQ water. HCR amplifiers B1 with AlexaFluor 647 as fluorophore were procured from Molecular Instruments, Inc. The specificity of the HCR reaction was meticulously validated through probe-negative staining.

Whole *L. stagnalis* embryos were carefully extracted from the egg capsules and fixed in 4% paraformaldehyde for 2 hours at room temperature. Subsequently, the samples underwent three washes with phosphate buffer (PBS) and were gradually dehydrated and stored in 100% methanol at –20°C. Preceding the HCR in situ hybridization (ISH) experiments, the samples were rehydrated in PBS through 10-minute steps. The Molecular Instruments HCR ISH protocol designed for whole- mount sea urchin embryos (Choi et al., 2018) with minor modifications was applied. Post HCR-ISH, selected samples underwent cryosectioning and were labeled with antibodies, as detailed below. The whole mount preparations were immersed in 2,2’-thiodiethanol and prepared for confocal scanning microscopy.

### 4.6 Western Blot Analysis

For Western blot analysis, *L. stagnalis* larvae at the veliger stage (st. 22) and the body part without the shell and visceral complex of postmetamorphic snails (st. 29) were utilized. The *L. stagnalis* embryos were carefully removed from the eggs and washed gently in phosphate buffer saline from the egg mucus (PBS, pH 7.4). Tissue samples were promptly sonicated at 4°C in RIPA buffer (150 mM NaCl, 1.0% NP40, 0.5% sodium deoxycholate, 0.3% SDS, 50 mM Tris, pH 8.0) and then centrifuged at 12,000 g for 30 minutes at 4°C. Supernatants were employed for subsequent investigations. Protein concentrations were determined using the BCA Protein Quantification Kit (Abcam, Cambridge, UK, ab102536) following the manufacturer’s instructions. The cleared homogenates were boiled for 5 minutes with β-mercaptoethanol. Polyacrylamide gels (10%) were loaded with samples (30 μg of protein/well), and electrophoresis was carried out for 30 minutes at 100V and 90 minutes at 160V in Tris/glycine/SDS running buffer. Proteins were then transferred to a nitrocellulose membrane (75 minutes at 80V in a transfer buffer containing 0.3% Tris, 1.44% glycine, and 30% methanol). To confirm the success of the transfer, the membranes were stained with Ponseau S solution. Nonspecific binding was blocked by a 1-hour incubation of the membrane in blocking buffer (TBS-T, 5% powdered milk), and membranes were subsequently incubated overnight at 4°C in blocking buffer with anti-Sox2 antibodies (Abcam, Cambridge, UK, ab97959, polyclonal, rabbit, 1:2000). Following several washes in TNT buffer, the membranes were incubated with anti-rabbit peroxidase-conjugated IgG (Jackson Immunoresearch, Cambridge, UK, 111-035- 144, goat, 1:5000) for 2 hours at room temperature. After the final washing in the TNT buffer, the membranes were revealed using the ECL detection system (Amersham Biosciences, UK, RPN2108).

### 4.7 Whole Mount and Cryosections Immunostaining

*L. stagnalis* embryos at various developmental stages were extracted from the eggs and thoroughly washed in PBS. Subsequently, the samples underwent a 3-hour fixation in 4% paraformaldehyde in PBS, followed by additional PBS washes. For the preparation of cryostat sections, selected samples were immersed in 20% sucrose in PBS for 24 hours at 4°C and subsequently frozen at −40°C. Sections with a thickness of 20 µm were generated using the Leica CM1950 cryostat (Leica, Germany) and affixed to glass slides. Immunolabeling procedures were consistent for both whole-mount preparations and cryostat sections, as well as for certain samples atter HCR ISH and EdU incorporation. Preparations were initially washed in PBS and then incubated for 1.5 hours at room temperature in 1% bovine serum albumin in PBS. Subsequently, preparations were exposed to various combinations of antibodies: anti-mouse Sox2 antibody (Abcam, Cambridge, UK, ab97959, polyclonal, rabbit, dilution 1:1000), anti-α-tubulin antibody (Sigma-Aldrich, Munich, Germany, T-6793, monoclonal, mouse, dilution 1:2000), and anti-FMRFamide antibody (Immunostar, Hudson, USA, 20091, polyclonal, rabbit, dilution 1:1000), all diluted in PBS containing 0.1% Triton-X100 and 0.1% bovine serum albumin, overnight at 4°C. After the antibody incubation, the preparations underwent a washing step and were then incubated in a mixture of secondary antibodies: anti-rabbit Alexa 488-conjugated IgG (Invitrogen, Waltham, USA; A-11008, goat, 1:700), anti-rabbit Alexa 555-conjugated IgG (Invitrogen, Waltham, USA; A-21428, goat, 1:700), and anti-mouse Alexa 633-conjugated IgG (Invitrogen, Waltham, USA; A-21050, goat, 1:700), all diluted in PBS containing 0.1% Triton-X100 and 0.1% bovine serum albumin, for 2 hours at room temperature. Following a final washing step, nuclei were stained with DAPI and washed in PBS again. Whole-mount preparations were immersed in 90% glycerol and then mounted on slides, while cryosections were enclosed in a hydrophilic medium Mowiol (Sigma-Aldrich, Munich, Germany, 81381).

### 4.8 Cell Proliferation Assays

Cell proliferation assays were conducted employing 5-Ethynyl-2′-deoxyuridine (EdU; ThermoFisher Cat# C10337), a thymidine analog, and a rat antibody specific to phosphorylated histone H3 (Sigma-Aldrich, H9908). For EdU labeling, *L. stagnalis* larvae at various developmental stages, extracted from the eggs, were incubated in 200 μM EdU diluted in Lymnaea saline solution (prepared according to Mescheriakov, 1990) for durations of 5 and 30 minutes. Subsequently, larvae underwent two washes in Lymnaea saline and were fixed in 4% paraformaldehyde for 3 hours. Following fixation, some samples underwent HCR ISH or antibody staining, employing the techniques described earlier. Visualization of EdU incorporation was achieved using the Click-iT EdU Alexa Fluor 488 Imaging kit (ThermoFisher Cat# C10337). Immunolabeling with a rat anti-phospho-histone H3 antibody (polyclonal rat antibody against phosphorylated Ser28 histone H3, Sigma-Aldrich, H9908) was carried out following the previously outlined techniques, and staining was detected using anti-rat Alexa 555-conjugated IgG (ThermoFisher, Catalog # A-21434).

### 4.9 Microscopy and Image proceeding

Preparations were examined utilizing a Zeiss LSM-880 confocal microscope (Carl Zeiss, Jena, Germany) with the application of appropriate wavelength-filter configurations. Subsequent to image acquisition, processing, and analysis of confocal images were performed using ZEN software (Carl Zeiss, Jena, Germany) and FIJI software (http://fiji.sc/Fiji).

## 5 Author Contributions

Study concept and design — EGI, EEV and AIK; collection of biological material — SNV, AIK and EEV; SNV — technical support; data acquisition — AIK, EGI; sequencing and transcriptome assembly – GRG, OSK, EIS; data analysis and interpretation — EGI, AIK and EEV (confocal microscopy), ADF and EGI (western blotting, bioinformatics analysis); writing and editing of the manuscript — AIK, EIG and EEV; administrative and material support — EEV. All authors have read and agreed to the published version of the manuscript.

## 7 Informed Consent Statement

Not applicable

## 8 Data Availability Statement

Identified sequences of L. stagnalis genes (Ls-SoxB1, Ls-SoxB2, Ls-SoxE, and Ls-Pangolin) were submitted to GenBank with the IDs OR853093, OR853091, OR853092, OR853094 respectively.

## 9 Acknowledgments

The research was done using the equipment of the Core Centrum of the Institute of Developmental Biology RAS. The research was conducted under IDB RAS RP # 0088-2021-0020.

## 11 Institutional Review Board Statement

Not applicable

## References

Angerer, L. M., Newman, L. A., and Angerer, R. C. (2005). SoxB1 downregulation in vegetal lineages of sea urchin embryos is achieved by both transcriptional repression and selective protein turnover. Development 132, 999–1008. doi: 10.1242/dev.01650.

Anishchenko, E., Arnone, M. I., and D’Aniello, S. (2018). SoxB2 in sea urchin development: implications in neurogenesis, ciliogenesis and skeletal patterning. EvoDevo 9, 5. doi: 10.1186/s13227-018-0094-1.

Anisimov, A. P. (2005). Endopolyploidy as a morphogenetic factor of development. Cell Biol Int 29, 993–1004. doi: 10.1016/j.cellbi.2005.10.013.

Anisimov, A. P., and Kirsanova, I. A. (2002). [Somatic polyploidy in neurons of the gastropod mollusca. III. Mitosis and endomitosis in the postnatal development of neurons in the Succinea snail central nervous system]. Tsitologiia 44, 981–987.

Arendt, D. (2018). Animal Evolution: Convergent Nerve Cords? Curr Biol 28, R225–R227. doi: 10.1016/j.cub.2018.01.056.

Arendt, D., Tosches, M. A., and Marlow, H. (2016). From nerve net to nerve ring, nerve cord and brain — evolution of the nervous system. Nat Rev Neurosci 17, 61–72. doi: 10.1038/nrn.2015.15.

Bakken, T. E., Miller, J. A., Ding, S.-L., Sunkin, S. M., Smith, K. A., Ng, L., et al. (2016). A comprehensive transcriptional map of primate brain development. Nature 535, 367–375. doi: 10.1038/nature18637.

Beklemishev, V. N. (1958a). Grundlagen der vergleichenden Anatomie der Wirbellosen: Promorphologie. VEB Deutscher Verlag der Wissenschaften Available at: https://books.google.de/books?id=F8AuAQAAIAAJ.

Beklemishev, V. N. (1958b). [On the early evolution of the molluscs]. Zoologicheskii Zhurnal 37, 518–522.

Buescher, M., Hing, F. S., and Chia, W. (2002). Formation of neuroblasts in the embryonic central nervous system of *Drosophila melanogaster* is controlled by *SoxNeuro*. Development 129, 4193– 4203. doi: 10.1242/dev.129.18.4193.

Bullock, T., and Horridge, G. A. (1965). Structure and function in the nervous systems of invertebrates. San Francisco.

Buresi, A., Andouche, A., Navet, S., Bassaglia, Y., Bonnaud-Ponticelli, L., and Baratte, S. (2016). Nervous system development in cephalopods: How egg yolk-richness modifies the topology of the mediolateral patterning system. Dev Biol 415, 143–156. doi: 10.1016/j.ydbio.2016.04.027.

Buresi, A., Croll, R., Tiozzo, S., Bonnaud, L., and Baratte, S. (2014). Emergence of Sensory Structures in the Developing Epidermis in Sepia officinalis and Other Coleoid Cephalopods. The Journal of comparative neurology 522. doi: 10.1002/cne.23562.

Bylund, M., Andersson, E., Novitch, B. G., and Muhr, J. (2003). Vertebrate neurogenesis is counteracted by Sox1-3 activity. Nat Neurosci 6, 1162–1168. doi: 10.1038/nn1131.

Camacho, C., Coulouris, G., Avagyan, V., Ma, N., Papadopoulos, J., Bealer, K., et al. (2009). BLAST+: architecture and applications. BMC Bioinformatics 10, 421. doi: 10.1186/1471-2105-10-421.

Choi, H. M. T., Schwarzkopf, M., Fornace, M. E., Acharya, A., Artavanis, G., Stegmaier, J., et al. (2018). Third-generation *in situ* hybridization chain reaction: multiplexed, quantitative, sensitive, versatile, robust. Development 145, dev165753. doi: 10.1242/dev.165753.

Chrysostomou, E., Flici, H., Gornik, S. G., Salinas-Saavedra, M., Gahan, J. M., McMahon, E. T., et al. (2022). A cellular and molecular analysis of SoxB-driven neurogenesis in a cnidarian. eLife 11, e78793. doi: 10.7554/eLife.78793.

Ciccarelli, F. D., Doerks, T., Von Mering, C., Creevey, C. J., Snel, B., and Bork, P. (2006). Toward Automatic Reconstruction of a Highly Resolved Tree of Life. Science 311, 1283–1287. doi: 10.1126/science.1123061.

Croll, R. P. (2000). Insights into early molluscan neuronal development through studies of transmitter phenotypes in embryonic pond snails. Microsc Res Tech 49, 570–578. doi: 10.1002/1097-0029(20000615)49:6<570::AID-JEMT7>3.0.CO;2-Q.

Croll, R. P. (2009). Developing Nervous Systems in Molluscs: Navigating the Twists and Turns of a Complex Life Cycle. Brain Behav Evol 74, 164–176. doi: 10.1159/000258664.

Croll, R. P., and Voronezhskaya, E. E. (1996). Early elements in gastropod neurogenesis. Dev Biol 173, 344–347. doi: 10.1006/dbio.1996.0028.

Deryckere, A., and Seuntjens, E. (2018). The Cephalopod Large Brain Enigma: Are Conserved Mechanisms of Stem Cell Expansion the Key? Front Physiol 9. doi: 10.3389/fphys.2018.01160.

Deryckere, A., Styfhals, R., Elagoz, A. M., Maes, G. E., and Seuntjens, E. (2021). Identification of neural progenitor cells and their progeny reveals long distance migration in the developing octopus brain. Elife 10, e69161. doi: 10.7554/eLife.69161.

Dong, Z., Shi, C., Zhang, H., Dou, H., Cheng, F., Chen, G., et al. (2014). The characteristics of sox gene in Dugesia japonica. Gene 544, 177–183. doi: 10.1016/j.gene.2014.04.053.

Duruz, J., Sprecher, M., Kaldun, J. C., Al-Soudy, A.-S., Lischer, H. E. L., van Geest, G., et al. (2023). Molecular characterization of cell types in the squid Loligo vulgaris. Elife 12, e80670. doi: 10.7554/eLife.80670.

Faccioni-Heuser, M. C., Zancan, D. M., and Achaval, M. (2004). Monoamines in the pedal plexus of the land snail Megalobulimus oblongus (Gastropoda, Pulmonata). Braz J Med Biol Res 37, 1043– 1053. doi: 10.1590/s0100-879x2004000700014.

Feuda, R., and Peter, I. S. (2022). Homologous gene regulatory networks control development of apical organs and brains in Bilateria. Sci Adv 8, eabo2416. doi: 10.1126/sciadv.abo2416.

Focareta, L., and Cole, A. G. (2016). Analyses of Sox-B and Sox-E Family Genes in the Cephalopod Sepia officinalis: Revealing the Conserved and the Unusual. PLoS One 11, e0157821. doi: 10.1371/journal.pone.0157821.

Fuchs, J., Martindale, M. Q., and Hejnol, A. (2011). Gene expression in bryozoan larvae suggest a fundamental importance of pre-patterned blastemic cells in the bryozoan life-cycle. EvoDevo 2, 13. doi: 10.1186/2041-9139-2-13.

Garstang, W. (1928). The Origin and Evolution of Larval Forms. Rep British Assoc Adv Sci Sect. D., 77–98.

Hartenstein, V., and Stollewerk, A. (2015). The Evolution of Early Neurogenesis. Dev Cell 32, 390–407. doi: 10.1016/j.devcel.2015.02.004.

Holmberg, J., Hansson, E., Malewicz, M., Sandberg, M., Perlmann, T., Lendahl, U., et al. (2008). SoxB1 transcription factors and Notch signaling use distinct mechanisms to regulate proneural gene function and neural progenitor differentiation. Development 135, 1843–1851. doi: 10.1242/dev.020180.

Ivashkin, E., Khabarova, M. Y., Melnikova, V., Nezlin, L. P., Kharchenko, O., Voronezhskaya, E. E., et al. (2015). Serotonin Mediates Maternal Effects and Directs Developmental and Behavioral Changes in the Progeny of Snails. Cell Rep 12, 1144–1158. doi: 10.1016/j.celrep.2015.07.022.

Jacob, M. H. (1984). Neurogenesis in Aplysia californica resembles nervous system formation in vertebrates. J. Neurosci. 4, 1225–1239. doi: 10.1523/JNEUROSCI.04-05-01225.1984.

Kalyaanamoorthy, S., Minh, B. Q., Wong, T. K. F., Von Haeseler, A., and Jermiin, L. S. (2017). ModelFinder: fast model selection for accurate phylogenetic estimates. Nat Methods 14, 587–589. doi: 10.1038/nmeth.4285.

Kandel, E. R., Kriegstein, A., and Schacher, S. (1980). Development of the central nervous system of Aplysia in terms of the differentiation of its specific identifiable cells. Neurosci 5, 2033–2063.

Karnavas, T., Mandalos, N., Malas, S., and Remboutsika, E. (2013). SoxB, cell cycle and neurogenesis. Front Physiol 4, 298. doi: 10.3389/fphys.2013.00298.

Katoh, K., and Standley, D. M. (2013). MAFFT Multiple Sequence Alignment Software Version 7: Improvements in Performance and Usability. Mol Biol Evol 30, 772–780. doi: 10.1093/molbev/mst010.

Kerner, P., Simionato, E., Le Gouar, M., and Vervoort, M. (2009). Orthologs of key vertebrate neural genes are expressed during neurogenesis in the annelid Platynereis dumerilii. EvoDevo 11, 513–524. doi: 10.1111/j.1525-142X.2009.00359.x.

Kiefer, J. C. (2007). Back to basics: Sox genes. Dev Dyn 236, 2356–2366. doi: 10.1002/dvdy.21218.

Kirsanova, I. A., and Anisimov, A. P. (2000). [Somatic polyploidy in neurons from gastropod mollusks. I. Morphological characteristics of ganglia and neurons in the CNS of the snail Succinea lauta]. Tsitologiia 42, 733–739.

Kirsanova, I. A., and Anisimov, A. P. (2001). [Somatic polyploidy in neurons of the gastropod molluscs. II. Dynamics of DNA synthesis in the process of postnatal growth of CNS neurons in Succineid snail]. Tsitologiia 43, 437–445.

Kuehn, E., Clausen, D. S., Null, R. W., Metzger, B. M., Willis, A. D., and Özpolat, B. D. (2022). Segment number threshold determines juvenile onset of germline cluster expansion in *Platynereis dumerilii*. J Exp Zool Pt B 338, 225–240. doi: 10.1002/jez.b.23100.

Larsson, A. (2014). AliView: a fast and lightweight alignment viewer and editor for large datasets. Bioinformatics 30, 3276–3278. doi: 10.1093/bioinformatics/btu531.

Le Gouar, M., Guillou, A., and Vervoort, M. (2004). Expression of a SoxB and a Wnt2/13 gene during the development of the mollusc Patella vulgata. Dev Genes Evol 214, 250–256. doi: 10.1007/s00427-004-0399-z.

Lindberg, D. R. (1988). “Heterochrony in Gastropods,” in Heterochrony in Evolution Topics in Geobiology., ed. M. L. McKinney (Boston, MA: Springer US), 197–216. doi: 10.1007/978-1-4899-0795-0_11.

Magie, C. R., Pang, K., and Martindale, M. Q. (2005). Genomic inventory and expression of Sox and Fox genes in the cnidarian Nematostella vectensis. Dev Genes Evol 215, 618–630. doi: 10.1007/s00427-005-0022-y.

Marlow, H., Tosches, M. A., Tomer, R., Steinmetz, P. R., Lauri, A., Larsson, T., et al. (2014). Larval body patterning and apical organs are conserved in animal evolution. BMC Biol 12, 7. doi: 10.1186/1741-7007-12-7.

Martín-Durán, J. M., Pang, K., Børve, A., Lê, H. S., Furu, A., Cannon, J. T., et al. (2018). Convergent evolution of bilaterian nerve cords. Nature 553, 45–50. doi: 10.1038/nature25030.

Martínez-Cerdeño, V., and Noctor, S. C. (2018). Neural Progenitor Cell Terminology. Front Neuroanat 12, 104. doi: 10.3389/fnana.2018.00104.

Masui, S., Nakatake, Y., Toyooka, Y., Shimosato, D., Yagi, R., Takahashi, K., et al. (2007). Pluripotency governed by Sox2 via regulation of Oct3/4 expression in mouse embryonic stem cells. Nat Cell Biol 9, 625–635. doi: 10.1038/ncb1589.

Meshcheryakov, V. N. (1990). “The Common Pond Snail Lymnaea stagnalis,” in Animal Species for Developmental Studies: Volume 1 Invertebrates, eds. T. A. Dettlaff and S. G. Vassetzky (Boston, MA: Springer US), 69–132. doi: 10.1007/978-1-4613-0503-3_5.

Monjo, F., and Romero, R. (2015). Embryonic development of the nervous system in the planarian Schmidtea polychroa. Dev Biol 397, 305–319. doi: 10.1016/j.ydbio.2014.10.021.

Moroz, L. L. (2009). On the Independent Origins of Complex Brains and Neurons. Brain Behav Evol 74, 177–190. doi: 10.1159/000258665.

Moroz, L. L. (2015). Biodiversity Meets Neuroscience: From the Sequencing Ship (Ship-Seq) to Deciphering Parallel Evolution of Neural Systems in Omic’s Era. *Integr. Comp. Biol.*, icv084. doi: 10.1093/icb/icv084.

Moroz, L. L. (2021). Multiple Origins of Neurons From Secretory Cells. Front Cell Dev Biol 9, 669087. doi: 10.3389/fcell.2021.669087.

Nezlin, L. P., and Voronezhskaya, E. E. (2017). Early peripheral sensory neurons in the development of trochozoan animals. Russ J Dev Biol 48, 130–143. doi: 10.1134/S1062360417020060.

Nguyen, L.-T., Schmidt, H. A., Von Haeseler, A., and Minh, B. Q. (2015). IQ-TREE: A Fast and Effective Stochastic Algorithm for Estimating Maximum-Likelihood Phylogenies. Mol Bio Evol 32, 268–274. doi: 10.1093/molbev/msu300.

Nielsen, C. (2012). Animal evolution: interrelationships of the living phyla. Oxford University Press.

Okuda, Y., Ogura, E., Kondoh, H., and Kamachi, Y. (2010). B1 SOX coordinate cell specification with patterning and morphogenesis in the early zebrafish embryo. PLoS Genet 6, e1000936. doi: 10.1371/journal.pgen.1000936.

Page, L. R. (1992). New Interpretation of a Nudibranch Central Nervous System Based on Ultrastructural Analysis of Neurodevelopment in Melibe leonina. I. Cerebral and Visceral Loop Ganglia. Biol Bull 182, 348–365. doi: 10.2307/1542255.

Panayi, H., Panayiotou, E., Orford, M., Genethliou, N., Mean, R., Lapathitis, G., et al. (2010). Sox1 Is Required for the Specification of a Novel p2-Derived Interneuron Subtype in the Mouse Ventral Spinal Cord. J Neurosci 30, 12274–12280. doi: 10.1523/JNEUROSCI.2402-10.2010.

Pevny, L., and Placzek, M. (2005). SOX genes and neural progenitor identity. Curr Opin Neurobiol 15, 7–13. doi: 10.1016/j.conb.2005.01.016.

Phochanukul, N., and Russell, S. (2010). No backbone but lots of Sox: Invertebrate Sox genes. Int J Biochem Cell Biol 42, 453–464. doi: 10.1016/j.biocel.2009.06.013.

Richards, G. S., and Rentzsch, F. (2014). Transgenic analysis of a SoxB gene reveals neural progenitor cells in the cnidarian Nematostella vectensis. Development 141, 4681–4689. doi: 10.1242/dev.112029.

Richter, S., Stach, T., and Wanninger, A. (2015). “Perspective—Nervous System Development in Bilaterian Larvae: Testing the Concept of ‘Primary Larvae,’” in Structure and Evolution of Invertebrate Nervous Systems (Oxford University Press Oxford), 313–324. doi: 10.1093/acprof:oso/9780199682201.003.0025.

Rizzo, N. P., and Bejsovec, A. (2017). SoxNeuro and shavenbaby act cooperatively to shape denticles in the embryonic epidermis of Drosophila. Development, dev.150169. doi: 10.1242/dev.150169.

Ross, K. G., Molinaro, A. M., Romero, C., Dockter, B., Cable, K. L., Gonzalez, K., et al. (2018). SoxB1 Activity Regulates Sensory Neuron Regeneration, Maintenance, and Function in Planarians. Dev Cell 47, 331–347.e5. doi: 10.1016/j.devcel.2018.10.014.

Saito, K., Michon, F., Yamada, A., Inuzuka, H., Yamaguchi, S., Fukumoto, E., et al. (2020). Sox21 Regulates Anapc10 Expression and Determines the Fate of Ectodermal Organ. iScience 23, 101329. doi: 10.1016/j.isci.2020.101329.

Sakharov, D. A. (1976). “Nerve cell homologies in gastropods,” in Neurobiology of Invertebrates: Gastropoda Brain: International Symposium on Invertebrate Neurobiology, Tihany, Hungary, Sept. 8–12, 1975, ed. J. Salánki (Budapest: Acad. Kiado), 27–40.

Schmidt-Rhaesa, A., Harzsch, S., and Purschke, G. eds. (2015). Structure and Evolution of Invertebrate Nervous Systems. 1st ed. Oxford: Oxford University Press doi: 10.1093/acprof:oso/9780199682201.001.0001.

Schock, E. N., York, J. R., Li, A. P., Tu, A. Y., and LaBonne, C. (2023). SoxB1 transcription factors are essential for initiating and maintaining the neural plate border gene expression. doi: 10.1101/2023.09.28.560033.

Semmler, H., Chiodin, M., Bailly, X., Martinez, P., and Wanninger, A. (2010). Steps towards a centralized nervous system in basal bilaterians: Insights from neurogenesis of the acoel Symsagittifera roscoffensis. *Dev.*, Grow. Diff. 52, 701–713. doi: 10.1111/j.1440-169X.2010.01207.x.

Shigeno, S., Sasaki, T., and von Boletzky, S. (2010). “The origins of cephalopod body plans: a geometrical and developmental basis for the evolution of vertebrate-like organ systems,” in Cephalopods-Present and Past, eds. K. Tanabe, Y. Shigeta, T. Sasaki, and H. Hirano (Tokyo: Tokai University Press), 23–34.

Shinzato, C., Iguchi, A., Hayward, D. C., Technau, U., Ball, E. E., and Miller, D. J. (2008). Sox genes in the coral Acropora millepora: divergent expression patterns reflect differences in developmental mechanisms within the Anthozoa. BMC Evol Biol 8, 311. doi: 10.1186/1471-2148-8-311.

Simionato, E., Kerner, P., Dray, N., Le Gouar, M., Ledent, V., Arendt, D., et al. (2008). atonal- and achaete-scute-related genes in the annelid Platynereis dumerilii: insights into the evolution of neural basic-Helix-Loop-Helix genes. BMC Evol Biol 8, 170. doi: 10.1186/1471-2148-8-170.

Somel, M., Franz, H., Yan, Z., Lorenc, A., Guo, S., Giger, T., et al. (2009). Transcriptional neoteny in the human brain. Proceedings of the National Academy of Sciences 106, 5743–5748. doi: 10.1073/pnas.0900544106.

Sur, A., Renfro, A., Bergmann, P. J., and Meyer, N. P. (2020). Investigating cellular and molecular mechanisms of neurogenesis in Capitella teleta sheds light on the ancestor of Annelida. BMC Evol Biol 20, 84. doi: 10.1186/s12862-020-01636-1.

Tan, S., Huan, P., and Liu, B. (2022). Molluskan Dorsal–Ventral Patterning Relying on BMP2/4 and Chordin Provides Insights into Spiralian Development and Evolution. Mol Biol Evol 39, msab322. doi: 10.1093/molbev/msab322.

Trifinopoulos, J., Nguyen, L.-T., von Haeseler, A., and Minh, B. Q. (2016). W-IQ-TREE: a fast online phylogenetic tool for maximum likelihood analysis. Nucleic Acids Res 44, W232–W235. doi: 10.1093/nar/gkw256.

Vagvolgyi, J. (1967). On the Origin of Molluscs, The Coelom, and Coelomic Segmentation. Systematic Zoology 16, 153. doi: 10.2307/2411408.

Vidal, B., Santella, A., Serrano-Saiz, E., Bao, Z., Chuang, C.-F., and Hobert, O. (2015). C. elegans SoxB genes are dispensable for embryonic neurogenesis but required for terminal differentiation of specific neuron types. Development 142, 2464–2477. doi: 10.1242/dev.125740.

Voronezhskaya, E. E., and Croll, R. P. (2015). “Mollusca: Gastropoda,” in Structure and Evolution of Invertebrate Nervous Systems (Oxford University Press Oxford), 196–221. doi: 10.1093/acprof:oso/9780199682201.003.0020.

Voronezhskaya, E. E., and Elekes, K. (1996). Transient and sustained expression of FMRFamide-like immunoreactivity in the developing nervous system ofLymnaea stagnalis (mollusca, pulmonata). Cell Mol Neurobiol 16, 661–676. doi: 10.1007/BF02151903.

Wei, Z., Angerer, R. C., and Angerer, L. M. (2011). Direct development of neurons within foregut endoderm of sea urchin embryos. Proceedings of the National Academy of Sciences 108, 9143–9147. doi: 10.1073/pnas.1018513108.

Wilson, M. J., and Dearden, P. K. (2008). Evolution of the insect Sox genes. BMC Evol Biol 8, 120. doi: 10.1186/1471-2148-8-120.

Wyeth, R. C., and Croll, R. P. (2011). Peripheral sensory cells in the cephalic sensory organs of Lymnaea stagnalis. Journal of Comparative Neurology 519, 1894–1913. doi: 10.1002/cne.22607.

