## Supplementary material for "Expanded Expression of Pro-Neurogenic Factor SoxB1 during Larval Development of Gastropod *Lymnaea stagnalis* Suggests Preadaptation to Prolonged Neurogenesis in Mollusca": Supplimental Table 1

### Supplementary file 1

**Table S1. Accession numbers of protein sequences used for phylogenetic tree construction**

| **NAME** | **NCBI accession number** | **NAME** | **NCBI accession number** |
| --- | --- | --- | --- |
| *Ac-SoxB1* | XP 005093681.1 | *Ls-Pangolin* | OR853094 |
| *Ac-SoxB2* | XP 005108229.1 | *Ls-SoxB1* | OR853093 |
| *Ac-SoxD* | XP 035824396.1 | *Ls-SoxB2* | OR853091 |
| *Ac-SoxE* | XP 005102100.1 | *Ls-SoxE* | OR853092 |
| *Ac-SoxF* | XP 005107482.1 | *Mm-Sox1* | BAC75667.1 |
| *Ce-SoxB* | NP 001335567.1 | *Mm-Sox2* | AAH57574.1 |
| *Ce-SoxB1* | NP 741836.1 | *Mm-Sox3* | AAL40744.1 |
| *Ce-SoxB2* | NP 510439.1 | *Mm-Sox4* | NP 033264.2 |
| *Ce-SoxC* | NP 740846.1 | *Mm-Sox5* | BAA32567.1 |
| *Ce-SoxD* | NP 001368074.1 | *Mm-Sox6* | AAC52263.1 |
| *Cg-SoxB1* | XP 011455662.1 | *Mm-Sox7* | NP 035576.1 |
| *Cg-SoxB2* | XP 011433975.1 | *Mm-Sox8* | NP 035577.1 |
| *Cg-SoxC* | XP 011445203.1 | *Mm-Sox9* | NP 035578.3 |
| *Cg-SoxD* | XP 011425377.1 | *Mm-Sox10* | NP 035567.1 |
| *Cg-SoxE* | AFK88538.1 | *Mm-Sox11* | NP 033260.4 |
| *Cg-SoxF* | XP 011448074.2 | *Mm-Sox12* | NP 035568.1 |
| *Cg-SoxH* | XP 011415859.3 | *Mm-Sox13* | NP 035569.2 |
| *Ct-Pangolin* | ELU10028.1 | *Mm-Sox14* | NP 033264.2 |
| *Ct-SoxB1* | AST23029.1 | *Mm-Sox17* | NP 001276393.1 |
| *Ct-SoxB2* | AST23030.1 | *Mm-Sox18* | NP 033262.2 |
| *Ct-SoxC* | ELT98138.1 | *Mm-Sox21* | NP 808421.1 |
| *Ct-SoxD* | ELT87450.1 | *Mm-Sox30* | AAF99391.1 |
| *Ct-SoxE* | ELU17008.1 | *Mm-Tcf7* | AAI45344.1 |
| *Ct-SoxF* | ELU06459.1 | *Pd-SoxB1* | ANS60443.1 |
| *Dm-Sox21b* | NP 001261828.1 | *Pd-SoxB2* | CAY12631.1 |
| *Dm-Dichaete* | NP 001261830.1 | *Pd-SoxC* | CAY12635.1 |
| *Dm-Pangolin* | NP 726528.2 |  |  |
| *Dm-Sox100B* | NP 651839.1 |  |  |
| *Dm-Sox102F* | NP 001014695.1 |  |  |
| *Dm-Sox14* | CAB64387.1 |  |  |
| *Dm-Sox15* | NP 523739.2 |  |  |
| *Dm-Sox21a* | NP 648694.1 |  |  |
| *Dm-Sox21b* | NP 001261828.1 |  |  |
| *Dm-SoxN* | CAB64386.1 |  |  |
| *La-Pangolin* | XP 013385963.1 |  |  |
| *La-SoxE* | XP 013381128.1 |  |  |
| *Lg-SoxB2* | XP 009044404.1 |  |  |
| *Lg-SoxH* | XP 009050874.1 |  |  |

Species abbreviations: Ac - *Aplysia californica*; Ce - *Caenorhabditis elegans*; Cg - *Crassostrea gigas*; Ct - *Capitella teleta*; Dm - *Drosophila melanogaster*; La - *Lingula anatina*; Lg - *Lottia gigantea*; Ls - *Lymnaea stagnalis*; Mm - *Mus musculus*; Pd - *Platynereis dumerilii*.
